# Myotendinous junctions form dynamic three-dimensional architectures during development and growth

**DOI:** 10.64898/2026.10.08.757742

**Authors:** Yuki Yamasaki, Yutaro Kawa, Ramu Sagasaki, Shunki Yashiro, Joonseop Shin, Mizuki Mishima, Kohei Arakawa, Hayate Suzuki, Seiya Mizuno, Kanon Otsuka, Tadayoshi Hayata, Chisa Shukunami, Masafumi Inui

## Abstract

The myotendinous junction (MTJ) is a specialized extracellular matrix-rich interface that transmits contractile force generated by skeletal muscle to tendon, but how the MTJ is assembled as developing skeletal muscle connects to tendon, and how the mature tissue-level architecture subsequently emerges, remain incompletely understood. Here, we generate a Col22a1-EGFP knock-in mouse that enables direct visualization and quantitative analysis of MTJs from embryonic development to postnatal growth. Tissue-clearing and three-dimensional imaging reveal extensive variation in MTJ morphology and spatial distribution among skeletal muscles and between myofiber types. During embryonic MTJ formation, COL22A1-EGFP undergoes a striking transition from intracellular puncta within developing myofibers to extracellular accumulation at myofiber tips, associated with polarized Golgi organization. We further identify complex tendon protrusions that emerge during postnatal growth and connect tendon bodies with subsets of MTJs. These interfaces contain morphologically distinct Scx-lineage cells that express Col22a1, identifying a specialized tendon-associated population at the MTJ. Finally, muscle unloading and overload differentially alter COL22A1-EGFP signal intensity and area, respectively, demonstrating that the reporter provides a quantitative readout of MTJ responses to altered mechanical conditions. Thus, Col22a1-EGFP reveals previously unrecognized spatial, developmental, and adaptive features of the muscle–tendon interface and provides an experimental platform for investigating MTJ biology in vivo.

## Introduction

The myotendinous junction (MTJ) is a transition tissue between skeletal muscle fibers and tendon extracellular matrix (ECM) that transmits the contractile force generated in muscle to tendon (Charvet et al., 2012; Mackey, 2024). The MTJ has specialized subcellular structure in which sarcolemma (myofiber membrane) at the tip of a myofiber and tendon collagen form interdigitations to increase the interface area (Charvet et al., 2012; Mair and Tomé, 1972). These structures have high adaptation capacity to mechanical loading, so that the depth of interdigitation decreases with muscle disuse and increases with exercise (Curzi et al., 2016; Kojima et al., 2008; Rocha et al., 2021; Sierra et al., 2018; Tidball, 1992). As MTJ is susceptible to injury and its regenerative potential is limited, it is considered as the weakest link in force transmission in locomotive organs (Edouard et al., 2023; Højfeldt et al., 2025; Jakobsen and Krogsgaard, 2021).

Gene expression and protein localization of MTJ are distinct from those in other parts of skeletal muscle to support its specialized structure and role in force transmission. Molecular characteristics of MTJ include ECM composition, transmembrane proteins and intracellular complexes of myofiber at the MTJ such as the enrichment of Laminin α2, Integrin α7, Dystrophin, Talin and Desmin (Law and Tidball, 1993; Mayer et al., 1997; Pedrosa-Domellöf et al., 2000; Shimizu et al., 1989; Tidball, 1992; Tidball et al., 1986). Recent transcriptome analysis at the single-nucleus level identified specific gene expression patterns of myonuclei close to MTJ region (Kim et al., 2020; Petrany et al., 2020; Santos et al., 2020), and proteome analyses identified several proteins specifically localized to MTJ (Karlsen et al., 2022; Schmidt et al., 2024). Among these molecular markers, the most distinct signature of MTJ is the presence of Type XXII collagen (COL22) (Koch et al., 2004).

Despite the increasing recognition of MTJ biology, the 3D organization of MTJ within intact tissues and how this tissue-level architecture is formed remains largely unexplored. In particular, it remains unknown how MTJs are spatially distributed across different muscle types, how MTJ architectures change during postnatal growth, and how MTJ structures are assembled during embryonic muscle-tendon interaction. In this study, to address this issue, we generated a Col22a1-EGFP knock-in mouse in which endogenous COL22A1 protein is fluorescently labeled in vivo.

Col22a1 belongs to fibril-associated collagen with interrupted triple helices (FACIT) type collagen family. The COL22A1 protein has a signal peptide, von Willebrand factor A (VWFA) domain, thrombospondin N-terminal-like domain (TSPN) and 5 collagenous domains with interruptions (Koch et al., 2004). It is expressed in several tissues in mammals, including skeletal muscle, heart, cartilage and skin (Feng et al., 2019; Koch et al., 2004). In skeletal muscle tissue, *Col22a1* mRNA is specifically expressed in myonuclei of MTJ (Kim et al., 2020; Petrany et al., 2020; Santos et al., 2020), and the protein localizes sharply at the interface of myofiber and tendon (Koch et al., 2004). As such, COL22A1 has been widely used as a standard MTJ marker from fish to humans (Charvet et al., 2013; Hoegsbjerg et al., 2025; Malbouyres et al., 2022). In addition to the specific expression at the adult MTJ, the earliest COL22A1 expression is reported in a decellularized mouse embryo at embryonic day 13.5-14.5 (E13.5-14.5) (Leng et al., 2025; Lipp et al., 2023), after the expected stage of muscle-tendon interaction (Huang et al., 2015; Ono et al., 2023; Sagasaki et al., 2026). Moreover, eQTL variants in *COL22A1* are associated with muscle injury in athletes (Miyamoto-Mikami et al., 2020) and in rat MTJ COL22A1 increased upon training after immobilization (Rocha-Braga et al., 2025), indicating that the expression of Col22a1 might not be merely an anatomical marker, but could be extended to a functional indicator of MTJ.

Combined with tissue clearing, light-sheet imaging, and developmental analyses, this model enabled visualization of MTJ architecture across multiple spatial and temporal scales. Using this approach, we demonstrate that MTJ is not a simple planar attachment structure but rather a dynamically remodeled and spatially heterogeneous 3D force-transmission architecture. We further identify specialized MTJ-associated tendon cells, developmental transitions in COL22A1 localization, and postnatal remodeling of tendon-MTJ topology.

## Materials and Methods

### Generation of Col22a1-EGFP mouse

We selected a sequence (5’-GCACACTTTTGCAACCTGCA -3’) in the 3rd exon of *Col22a1* as the crRNA. We purchased synthetic crRNA containing this target in sequence from IDT (Iowa, US). Donor DNA was constructed as follows: genomic regions of 1,069 bp (1,061 bp of intron 2 and 8 bp of exon 3) upstream and 1,200 bp downstream (559 bp of exon 3 and 641 bp of intron 3) were used as 5’ and 3’ homology arms, respectively. EGFP coding sequence flanked by GS linker sequences was cloned between the 5’ and 3’homology arms (Fig. 1A, S1A). The isolation and preparation of the donor DNA vector, crRNA, tracrRNA, and Cas9, microinjection into C57BL/6J mouse zygotes, and embryo transfer were performed as previously described (Tanimoto et al., 2022).

**Figure 1.**
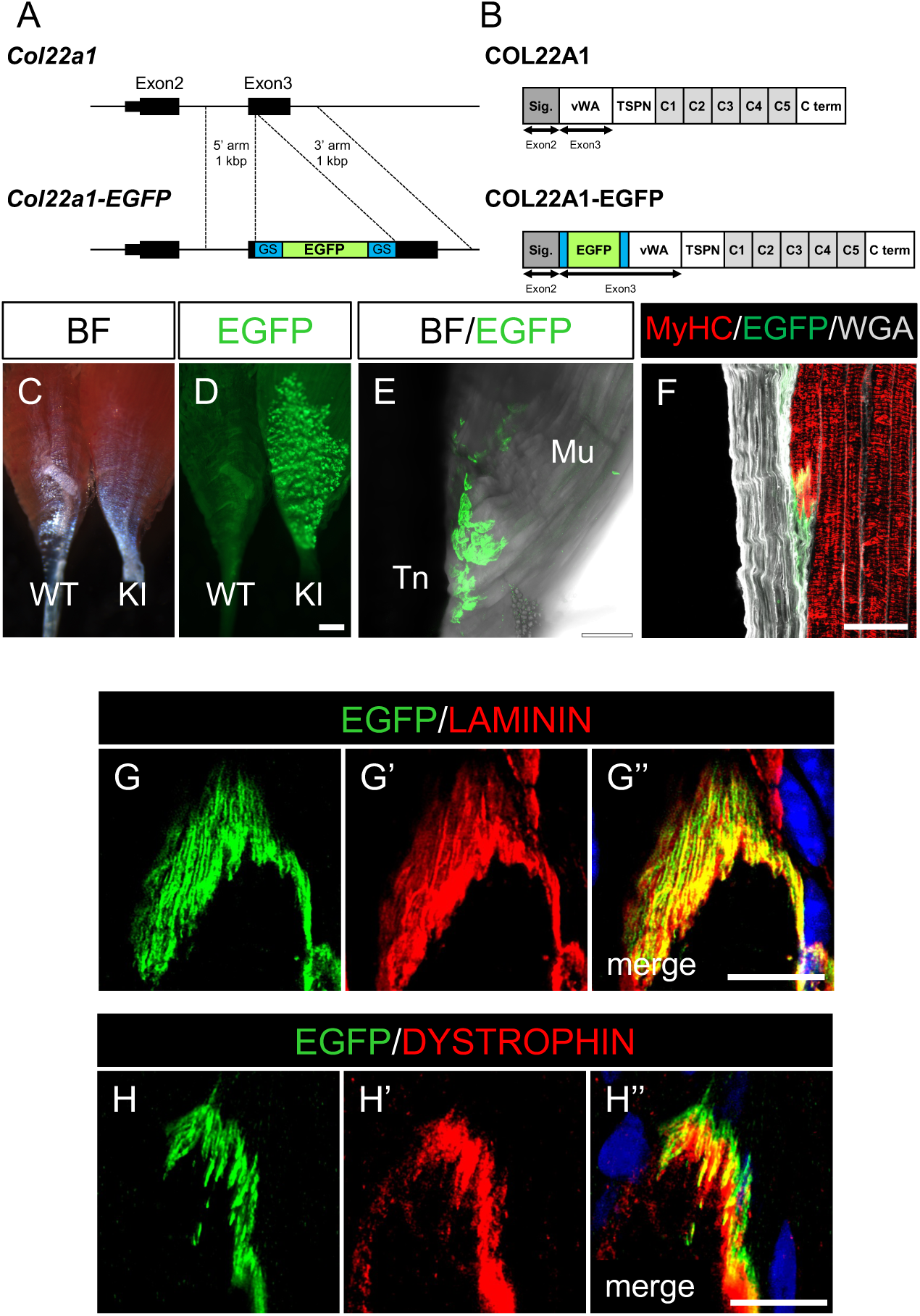
Generation of Col22a1-EGFP mice. (A) A schematic drawing of Col22a1-EGFP knock-in design. GS, GS linker sequence (B) A schematic drawing of COL22A1-EGFP fusion protein. Sig, signal sequence; VWFA, von Willebrand factor A-like domain; TSPN, N-terminal thrombospondin-like domain; C1-5, collagen repeat domain 1-5; C term, c-terminal domain. (C, D) Stereomicroscope image of freshly isolated SOL muscles from wild-type (WT) and Col22a1-EGFP knock-in (KI) mice. BF, bright field. Scale bar: 1 mm (E) Confocal microscope image of freshly isolated SOL muscle from Col22a1-EGFP knock-in mouse. Tn, tendon; Mu, muscle. Scale bar: 50 μm (F) Immunofluorescence image of Extensor Digitorum Longus (EDL) muscle section of Col22a1-EGFP knock-in mouse. MyHC, myosin heavy chain; WGA, wheat germ agglutinin, Scale bar: 50 μm (G, H) Immunofluorescence image of EDL muscle section of Col22a1-EGFP knock-in mouse. Blue signals are nuclei stained with DAPI. Scale bar: 10 μm

### Mice

*ScxCreERT2* transgenic (Tg) and *ScxGFP* Tg mouse strains have been described previously (Sugimoto et al., 2013; Yimiti et al., 2025). *Rosa26-LSL-tdTomato* (Ai9) strain (Madisen et al., 2010) was purchased from Jackson Laboratory (strain ID 7909). C57BL/6 and ICR mice were purchased from Sankyo lab-service (Tokyo, Japan). *ScxCreERT2;Rosa26-LSL-tdTomato* mice were subjected to intraperitoneal tamoxifen injections. Each animal received a total of five injections on alternating days, each containing approximately 75 mg/kg body weight of tamoxifen (Sigma-Aldrich, St. Louis, USA) and dissected 5 days after the last tamoxifen administration. All animal experiments were approved by the animal experiment committees of Meiji University (approval No. MUIACUC2022-04).

### Dissection and surgery models

For adult muscles, 8–12-week-old mice were sacrificed by cervical dislocation, skinned and skeletal muscles were isolated with dissection scissors and forceps. For embryo sampling, pregnant female mice were sacrificed by cervical dislocation, skinned and embryos were isolated from the uterus. Isolated muscles and embryos were imaged with fluorescence stereomicroscope (MZ-10F, Leica, Wetzlar, Germany) and confocal microscope (Stellaris 8, Leica). For sciatic denervation, 8–12-week-old male mice were anesthetized, the skin posterior to the femur was incised, the sciatic nerves were exposed, and 1-2 mm of the nerve were removed by cutting two sites with microscissors. The incision was sutured, and the mice were sacrificed after 14 days. For tenotomy, 8–12-week-old male mice were anesthetized and incised at the distal end of the gastrocnemius muscle. Achilles’ tendon was exposed, and distal tendons of gastrocnemius and soleus muscle (SOL) were cut off by microscissors. The incision was sutured and the mice were sacrificed after 14 days.

### Immunofluorescence

Adult muscle tissues were immersed in 10% and 30% sucrose in PBS and embedded in optimal cutting temperature (OCT) compound (Sakura finetek, Tokyo, Japan) without a fixation process. Embryonic samples were fixed with 4% paraformaldehyde (PFA) for 3 minutes, immersed in 10% and 30% sucrose in PBS and embedded in OCT compound. Tissues were sectioned at 10-20 μm and subjected to immunofluorescence. Adult sections were washed in PBS to remove OCT compound, post-fixed in 4% PFA in PBS for 5 minutes. Sections were then blocked with 5% horse serum, 2% bovine serum albumin in PBS, incubated with primary antibody solution overnight at 4 °C. For myofiber type analysis, blocking was performed with M.O.M. blocking reagent (Vector, USA). After washing in PBST (PBS supplemented with 0.25% Tween 20), sections were incubated with a secondary antibody solution for 1 hour at room temperature in the dark. Following a wash in PBST, sections were mounted with a mounting medium. Embryo sections were washed in PBS to remove OCT compound and post-fixed in 4% PFA in PBS for 5 minutes, followed by a wash in PBS. Sections were then blocked with 2% skim milk in PBS for 1 hour at room temperature and incubated with primary antibody solution overnight at 4 °C. After washing in PBST, sections were incubated with a secondary antibody solution for 1 hour at room temperature in the dark. Following a wash in PBST, sections were mounted with a mounting medium. Images were obtained by a confocal microscope (Stellaris 8) with z-stack and maximum projection. α-Laminin (Sigma-Aldrich, L9393, 1:500), α-Dystrophin (Abcam, Cambridge, UK, ab15277, 1:250), α-Col22a1 (Abcam, ab121846, 1:200), α-Type I collagen (SouthernBiotech, Birmingham, AL, USA, 1310-01, 1:1000), α-GFP (Abcam, ab13970, 1:500), α-Myosin heavy chain (Developmental Studies Hybridoma Bank (DSHB), Iowa City, IA, USA, MF20, 1:200, or Sigma-Aldrich, My-32, 1:500), α-Type I myosin heavy chain (DSHB, BA-D5, 1:50, 5.18 µg/mL), α-Type IIa myosin heavy chain (DSHB, SC-71, 1:10, 4.9 µg/mL), α-GM130 (BD Transduction, New Jersey, USA, #610823, 1:200), and α-Scleraxis (Kumagai et al., 2012) were used as primary antibodies. α-rabbit IgG Cy3, α-rabbit IgG-Alexa fluor 488, α-mouse IgG Cy3, α-chicken IgY Cy3, α-chicken IgY Cy5, or α-goat IgG-Alexa488 (Jackson Immunoresearch, West Grove, USA) were used as secondary antibodies. α-mouse IgG1 Alexa555 (Invitrogen, A21127, 1:250) and α-mouse IgM DyLight405 (Jackson Immunoresearch, 115-477-187, 1:250) were used to detect α-Type I myosin heavy chain and α-Type IIa myosin heavy chain, respectively. Slides were coverslipped with Fluoroshield Mounting Medium with DAPI (ImmunoBioScience Corp., WA, USA).

### Quantification of endogenous EGFP signal

To quantify the signal intensity and EGFP positive area of MTJs, images of MTJs were obtained as described in the immunofluorescence section, but antibodies against COL22A1 or EGFP were not included and fluorescence signal from COL22A1-EGFP protein was directly detected using a 488 nm laser. Images were then processed using ImageJ software. COL22A1-EGFP fluorescence at the MTJs was analyzed using maximum-intensity projection images generated from z-stack images. Images were acquired using a confocal microscope (Stellaris 8) with a 40x objective. Z-stack images were obtained with a total stack size of 6 µm. For each sample, at least two sections were imaged, corresponding to approximately 30 or more MTJs. Maximum-intensity projection images were generated using LAS X software. Individual MTJ regions were manually selected using the “Polygon Selection” tool in ImageJ/Fiji. To quantify the EGFP-positive area, isolated MTJ images were converted to 8-bit grayscale images, and EGFP-positive pixels were segmented using a fixed intensity range of 13–255. The resulting EGFP-positive region was converted into a binary mask, and the area of the signal-positive pixels was measured using ImageJ/Fiji. The EGFP-positive area was measured for each MTJ using the “Measure” tool. To quantify EGFP fluorescence intensity, the pixel-intensity distribution of each isolated MTJ image was obtained using the “Histogram” tool in ImageJ/Fiji. Histogram data were exported as CSV files, and only pixels with intensity values between 13 and 255 were used for analysis. The mean fluorescence intensity of each MTJ was calculated as a weighted mean using the following formula:

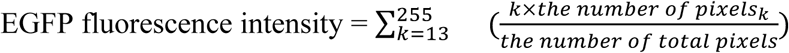

For both quantifications, at least 30 MTJs were analyzed per animal, and quantifications from three animals were pooled and compared between conditions.

### Quantification of Peri-nuclear Golgi apparatus distribution

The position of the Golgi apparatus relative to each myonucleus was quantified using ImageJ/Fiji. Forelimb sections from P0 Col22a1-EGFP mice (n = 3) were immunostained with α-GM130, α-Laminin, and α-GFP antibodies and counterstained with DAPI as described above. Images were acquired with a 40x objective, and maximum-intensity projections of z-stack images were used for analysis. Four DAPI+ myonuclei closest to MTJ, judged by COL22A1-EGFP signal, were defined as “MTJ-proximal” nuclei and myonuclei located sixth or more distal from MTJ were defined as “MTJ-distant” nuclei. Only Golgi apparatus overlapping with a single myonucleus were included in the quantification and myonuclei and GM130-positive Golgi apparatus were manually annotated and cropped. Myonuclei manually annotated above were approximated with an ellipse by ImageJ ellipse tool and its major axis was defined. We considered this major axis of myonucleus as surrogate myofiber longitudinal axis to overcome local bending of myofibers. We calculated the angle of “Golgi apparatus-nuclear centroid line” to nuclear major axis as the parameter of Golgi apparatus distribution relative to myonuclei, defining 0° when two lines overlap on the MTJ side of the nucleus, and 180° on the opposite side. A vector along the nuclear major axis from the nuclear centroid to the edge of the nucleus toward the MTJ is defined as vector **a**. Another vector from the nuclear centroid to Golgi centroid is defined as vector **b**. The angle θ between **a** and **b** was calculated. At least 20 Golgi apparatus were examined for each animal, pooled and their distribution were visualized with rose plots.

### Fluorescent *in situ* hybridization

Hybridization chain reaction-based fluorescent in situ hybridization method ISH palette (Nepagene, Chiba, Japan) was performed according to the manufacturer’s instruction and to a previous report (Tsuneoka and Funato, 2020). Briefly, cryosection of embryonic and adult tissues were hybridized with 20-30 pairs of single strand DNA probes at 37 °C overnight. Samples were washed and reacted with hairpin DNA with fluorescent dye for 5 hours at room temperature in the dark, further washed with 0.25% PBST and PBS, stained with DAPI and cover-slipped with Fluoroshied mounting reagent. Images were obtained by a confocal microscope (Stellaris 8) with z-stack and maximum projection. The probe sequences used were shown in Supplementary Table1.

### Tissue clearing and whole-mount immunofluorescence

Tissue clearing with CUBIC reagents (Susaki et al., 2014; Susaki et al., 2020) was performed as described previously (Sagasaki et al., 2026). Briefly, adult muscle tissues and embryos were fixed with 4% PFA for 30 minutes at 4°C. Samples were washed with PBS, embryo samples were digested with 10 μg/mL Proteinase K at 37°C for 30 minutes and re-fixed with 4% PFA/0.2% glutaraldehyde at room temperature for 20 minutes. Embryo and adult samples were incubated with 50% CUBIC1/H_2O_ for 2-4 hours at 37°C followed by 100% CUBIC1 at 37°C overnight with gentle rotation. Samples were then washed twice with PBS for 10 minutes, blocked with 1% bovine serum albumin/5% horse serum in PBS (hereafter blocking solution) for 2-4 hours, and incubated with primary antibodies in blocking solution at 4°C for two overnights. Samples were washed 10 times with PBSTx (PBS supplemented with 0.1% Triton-X100) for 40 minutes, blocked with blocking solution for 2-4 hours and incubated with secondary antibodies at 4°C overnight. Samples were washed 10 times with PBSTx for 40 minutes and further cleared with CUBIC2. Samples were mounted on a glass-bottom dish with 2% low-melting agarose/CUBIC2 gel and images were obtained with a confocal microscope (Stellaris 8, light-sheet unit). 300-600 slices were processed with the LAS X program as 3D images and virtual cross sections. Primary antibodies used were α-GFP (Abcam, 1:1000), α-Type I Collagen (SouthernBiotech, 1:1000), α-Goat IgG-Alexa488 (#705-545-003, Jackson ImmunoResearch, 1:1000), and α-chicken IgY-Cy3 (Jackson ImmunoResearch, #703-165-155, 1:1000).

### 3D micro-computed tomography (µCT)

24-week-old male mice were sacrificed by cervical dislocation and femurs were isolated. Isolated femurs were fixed at 4°C in 100% ethanol and stored in 100% ethanol until scanning. 3D micro-computed tomography (µCT) images were obtained using the ScanXmate-E090 (Comscantechno, Yokohama, Japan) at 90 kV and 90 μA, with a resolution of 10 μm/pixel. Images were reconstructed using coneCTexpress software (Voxel Works, Tokyo, Japan), and bone density was corrected using a phantom (Ratoc System Engineering, Tokyo, Japan). Trabecular bone morphometric parameters, including bone volume/tissue volume (BV/TV), were calculated on the distal femur using TRI/3D-BON software (Ratoc System Engineering, Tokyo, Japan). The region of interest (ROI) was set starting at 500 μm proximal to the growth plate and extending 1000 μm proximally. Statistical comparisons between two independent groups were performed using Welch’s t-test, which does not assume equal variances.

## Results

### Generation of Col22a1-EGFP mouse

To generate a reporter mouse that allows us to visualize the MTJ dynamics, we selected Col22a1 as the target molecule (Kim et al., 2020; Koch et al., 2004). EGFP coding sequence flanked by GS-linker sequences was knocked into the third exon of *Col22a1*, expecting that COL22A1 protein with EGFP fused at its N-terminus would be secreted to MTJ (Fig. 1A, B). crRNA targeting the third exon of *Col22a1*, tracrRNA and Cas9 protein together with knock-in targeting vector were microinjected into C57BL/6 fertilized eggs and 2-cell embryos were transferred to pseudo-pregnant ICR females. We obtained pups with EGFP sequence knocked into the expected locus (Fig. S1A, B), and Col22a1-EGFP line was established from a founder male. Col22a1-EGFP allele was transmitted at the expected Mendelian ratio and homozygous Col22a1-EGFP mice were indistinguishable from their wild-type littermates (data not shown). Loss of *Col22a1* function is reported to result in decreased trabecular bone mass (Zhao et al., 2021), but homozygous Col22a1-EGFP knock-in mice did not show an apparent reduction of trabecular bone volume compared to heterozygous mice (Fig. S1C, D), implying that EGFP-fusion at N-terminus does not grossly impair Col22a1 function. Nevertheless, we analyzed only heterozygous reporter mice throughout this study to minimize any potential effects of the EGFP fusion.

In freshly isolated soleus (SOL) muscle of Col22a1-EGFP knock-in mouse, discrete EGFP fluorescence signals were detected with stereo microscopy (Fig. 1C, D). Confocal microscope imaging (wholemount and section) confirmed that the EGFP fluorescence was detected at the interface of skeletal muscle and tendon tissue (Fig. 1E, F). COL22A1-EGFP fusion protein was detected at the distal MTJ of tibialis anterior (TA) muscle in a similar manner to the endogenous COL22A1 protein (Fig. S1E, F). These results indicated that our knock-in mouse expressed the EGFP reporter in myotendinous junction as designed. Moreover, the Col22a1-EGFP signal overlapped with that of extracellular matrix protein Laminin (Fig. 1G, G’, G’’) but was detected slightly outside of Dystrophin lining the sarcolemma (Fig. 1H, H’, H’’), indicating that the fusion protein is secreted to the extracellular space. These findings suggested that the Col22a1-EGFP reporter mouse faithfully recapitulates the localization of endogenous COL22A1 protein, enabling us to visualize MTJs at cell and tissue levels.

### Col22a1-EGFP revealed wide variations of shape and distribution of myotendinous junction among skeletal muscle types

Having established an MTJ reporter mouse, we next wanted to reveal 3D structure of individual MTJ as well as the variation of MTJs among muscle types (e.g. pennate muscle vs fusiform muscle) or myofiber types (slow fiber vs fast fiber).

EGFP signals of the distal end of the Extensor Digitorum Longus (EDL) muscle, a pennate muscle predominantly composed of fast-twitch-fibers, were observed as discrete spots distributed along the muscle-tendon interface that lies on the single side of the muscle (Fig. 2A, A’, yellow arrowhead). In tissue section, EGFP positive MTJs were aligned at the interface of tendons and muscle tissue as a thin layer (Fig. 2B). SOL, another pennate muscle rich in slow-twitch fibers, also demonstrated a discrete pattern, but with broader and brighter signals (Fig. 2C, C’, D), indicating that MTJ morphology and distribution are not uniform among muscles. Biceps, an example of fusiform muscle, showed EGFP signals evenly distributed (not biased to the single side) at its distal end (Fig. S2A, B). Precisely aligned EGFP signal was detected between the segments of rectus abdominis muscle and at the boundary of the diaphragm muscle and its central tendon (Fig. S2C-F). Extraocular muscle displayed a sharp pinhead-like EGFP signal at the distal end (Fig. S2G, H). Tongue and hamstring muscles also exhibited EGFP signals at their tendon interfaces with distinct patterns (Fig. S2I-L). These results demonstrated that MTJ distribution is highly diverse among muscle types.

**Figure 2.**
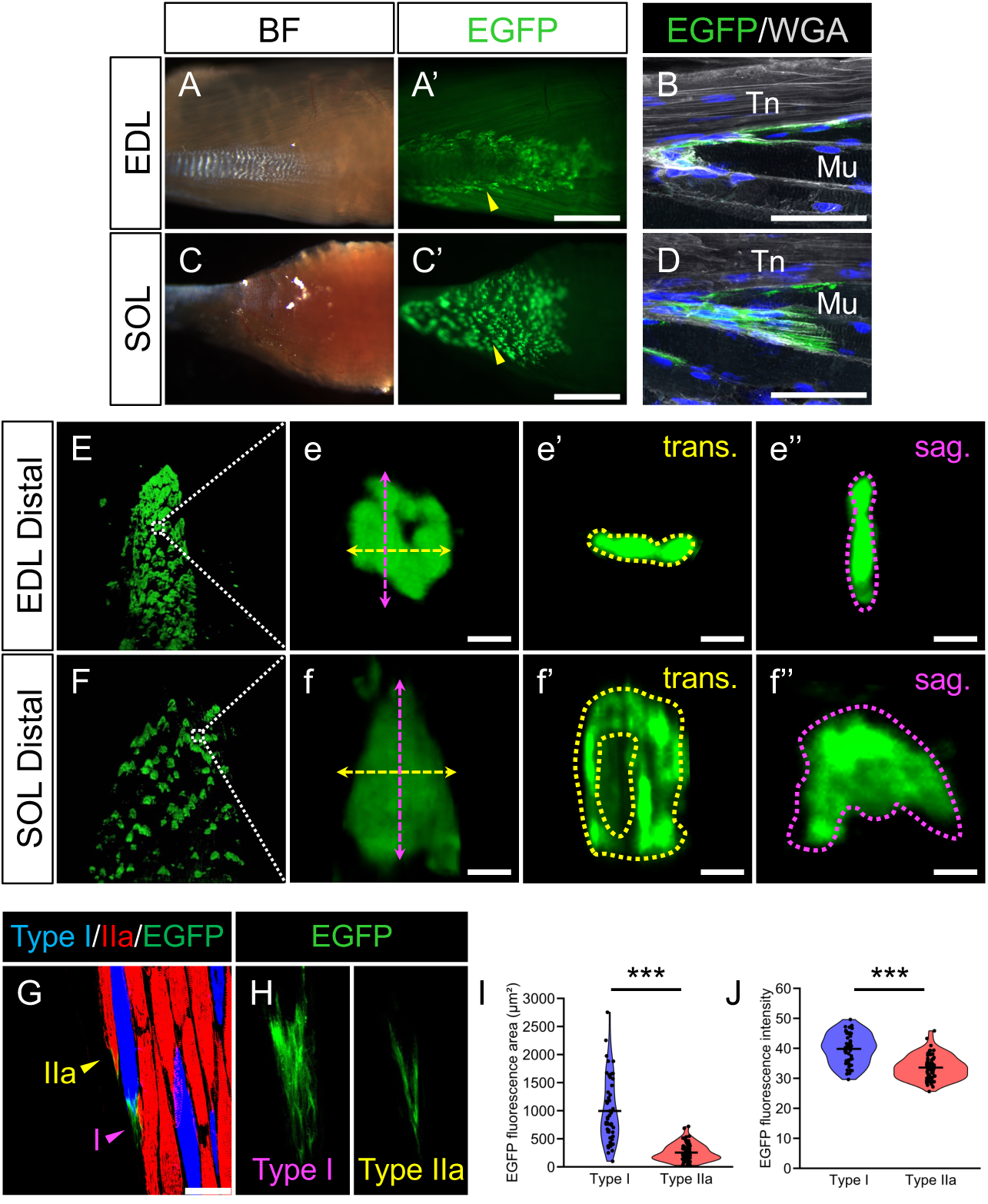
Col22a1-EGFP revealed variations in MTJ shape, size, and distribution among skeletal muscle types as well as myofiber types. (A, A’) Stereomicroscope images of EDL muscle of Col22a1-EGFP mice. (B) Immunofluorescence image of EDL muscle section of Col22a1-EGFP knock-in mouse. Blue signals are nuclei stained with DAPI. Tn, tendon; Mu, muscle. Scale bar: 50 μm (C, C’) Stereomicroscope images of SOL muscle of Col22a1-EGFP mice. (D) Immunofluorescence image of SOL muscle section of Col22a1-EGFP knock-in mouse. Blue signals are nuclei stained with DAPI. Scale bar: 50 μm (E, F) Light-sheet image of the distal end of EDL (E) and SOL (F) muscle of Col22a1-EGFP mice. Scale bar: 200 µm. (e) and (f) are the magnified views of the boxed areas shown in E and F Scale bar: 20 µm. e’/e’’ and f’/f’’ display a digital section of MTJ shown in e and f at dashed lines. Yellow lines (e’/f’) are cross-sections and magenta lines (e’’/f’’) are sagittal-sections along the proximal-distal axis of each muscle. (G, H) Immunofluorescence image of SOL muscle section of Col22a1-EGFP knock-in mouse. Blue signals are nuclei stained with DAPI. I, type I fiber; IIa, type IIA fiber. Scale bar: 50 μm (I, J) Quantification of COL22A1-EGFP positive area (I) and mean COL22A1-EGFP signal intensity (J) from immunofluorescence images. Statistical analyses were performed using Welch’s t-test, ***p<0.001

Tissue-clearing and light-sheet observation further revealed the variation of MTJ 3D structures between EDL and SOL muscles. While the Col22a1-EGFP signal of MTJs of EDL showed flat and scale-like morphology at the interface between myofiber and tendon (Fig. 2E, e-e’’’), those of SOL showed dome-shaped morphology capping the tip of myofibers (Fig. 2F, f-f’’’). Distribution of MTJs seems dense in EDL and sparse in SOL. Intriguingly, MTJ morphologies of EDL and SOL in neonates were not as distinct as those in adults, indicating that the MTJ morphology diversifies during the postnatal growth period (Fig. S5C, D).

Lastly, MTJs of type I and IIa fibers were compared in the sagittal section of SOL muscle (Fig. 2G, H). We found that COL22A1-EGFP positive area and signal intensity of type I fibers were significantly higher than those of type IIa fibers (Fig. 2I, J). Together, Col22a1-EGFP mouse revealed that the morphology and distribution patterns of MTJs in skeletal muscle tissues have striking variability, which is in part dynamically acquired postnatally.

### Col22a1-EGFP highlights muscle-tendon connection and intracellular MTJ formation dynamics during embryonic development

Next, we observed the initial step of MTJ formation. In developing embryos, COL22A1-EGFP signal was observed as early as E12.5 at immunostained section (Fig. 3A, E). Initially, COL22A1-EGFP signals were detected as scattered particle-like signals within myofibers (Fig. 3e, e’, yellow arrowheads), at the proximal, but not the distal end of forelimb muscles (Fig. 3E, S3). At E14.5, in addition to the intra-myofiber dot-like signal, extracellular COL22A1-EGFP signal at the tip of myofiber was first detected (Fig. 3B, F, f, f’, yellow brackets). At later stages, while the intra-myofiber signal gradually diminished, the extracellular signal accumulated progressively (Fig. 3C, D, G, H). These data suggest that the COL22A1-EGFP protein gradually shifted from intracellular to extracellular locations as the embryonic stage proceeded, revealing unprecedented protein dynamics of the MTJ formation process.

**Figure 3.**
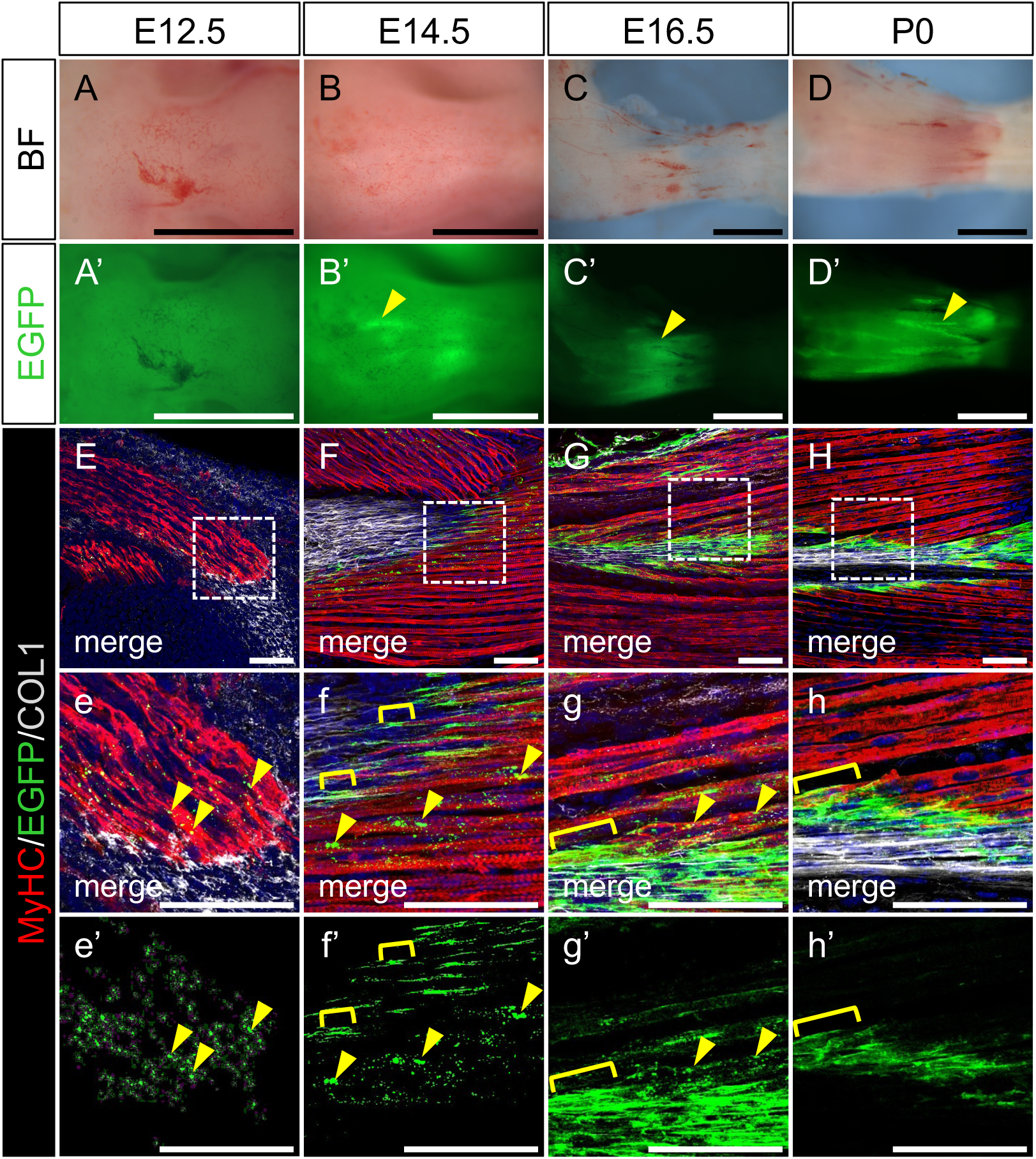
Embryonic expression of Col22a1-EGFP. (A-D, A’-D’) Stereomicroscope images of Col22a1-EGFP embryo forelimbs at E12.5 (A, A’), E14.5 (B, B’), E16.5 (C, C’), and P0 (D, D’). Yellow arrowheads indicate EGFP signals at the putative proximal end (A, A’) and distal end (B-D, B’-D’) of the extensor digitorum communis muscle. Scale bar: 1 mm. (E-H) Immunofluorescence image of Col22a1-EGFP embryonic forelimb sagittal sections at E12.5 (E), E14.5 (F), E16.5 (G), and P0 (H). (e)-(h), (e’)-(h’) are the magnified views of the boxed area shown in E-H. The yellow arrowheads indicate dot-like signals within the myofibers. The yellow brackets indicate putative extracellular signals at the tip of myofibers. Blue signals are nuclei stained with DAPI. Scale bar: 50 µm.

In contrast to the dynamic change of COL22A1-EGFP protein localization between E14.5 and E18.5 (Fig. 4A, B), *Col22a1* mRNA localization remained relatively unchanged (Fig. 4C, D). At both stages *Col22a1* mRNA transcribed in myonuclei close to the fiber tip was distributed toward the tip and central side of myofiber, likely by passive diffusion (Fig. 4C, D). These results imply mechanisms for the polarized secretion and/or retention of COL22A1 protein at MTJ. In line with this idea, we found that the peri-nuclear Golgi apparatus near myofiber tips were polarized toward the tips, whereas those distant from the tips showed no polarization (Fig. 4E-G). Altogether, our findings demonstrated that the Col22a1-EGFP is a unique reporter that enables us to visualize muscle-tendon interaction as well as MTJ formation dynamics. Muscle-tendon connection occurs at approximately E12.5-E13.5, coinciding with the earliest detection of COL22A1-EGFP protein. Muscle-tendon connection has been assessed by morphological/histological evaluation, while specific molecular markers have rarely been used (Ono et al., 2023; Tanji et al., 2023). The Col22a1-EGFP reporter could therefore serve as a surrogate marker of muscle-tendon interaction in embryos.

**Figure 4.**
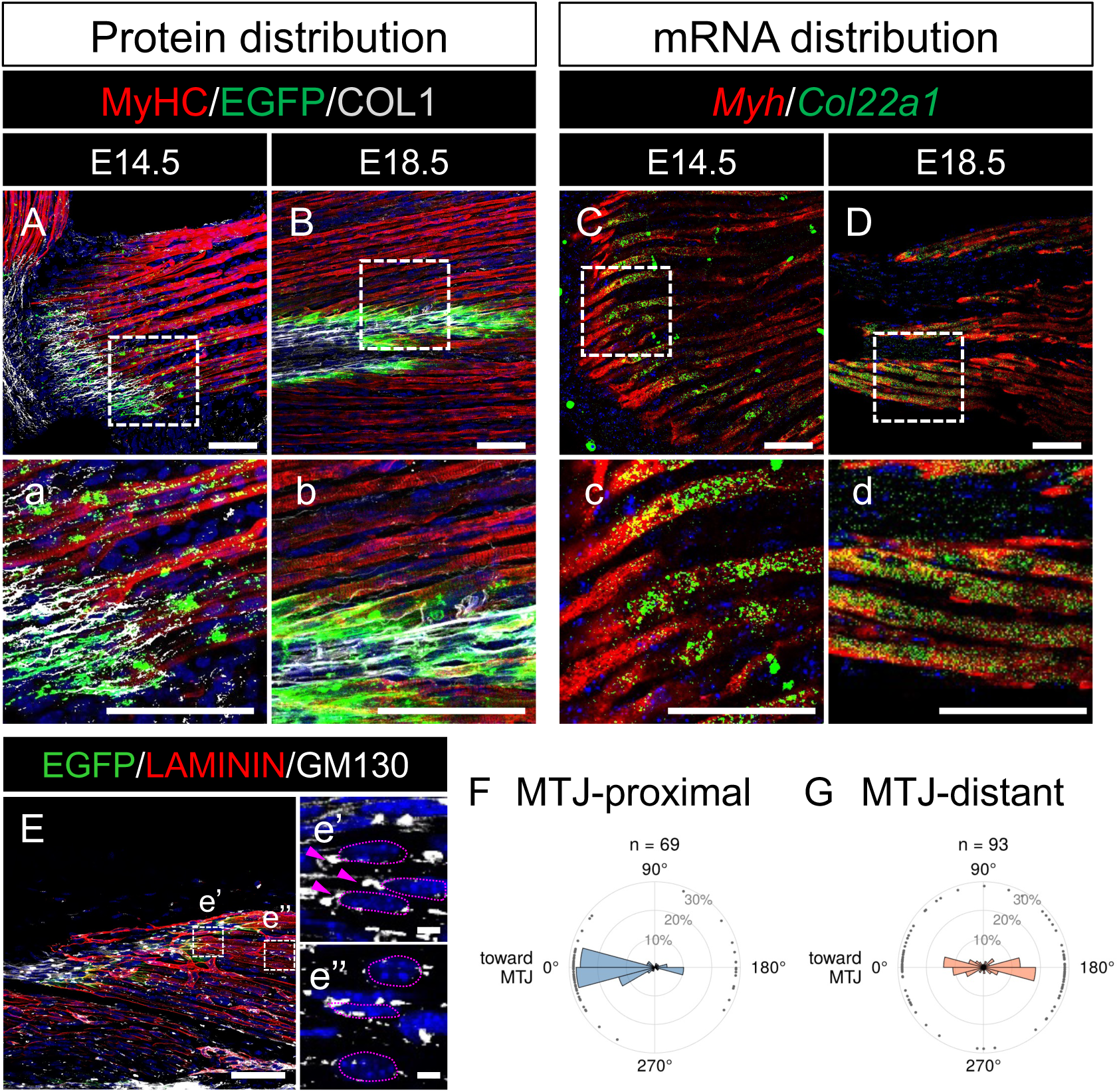
Comparison of COL22A1-EGFP protein and *Col22a1* mRNA distribution in embryonic myofibers. (A, B) Immunofluorescence images of Col22a1-EGFP embryo forelimbs at E14.5 (A) and E18.5 (B). (a) and (b) are the higher magnification views of boxed areas in A and B. Blue signals are nuclei stained with DAPI. Scale bar: 50 µm (C, D) Fluorescence ISH image of wild-type embryo forelimbs at E14.5 (C) and E18.5 (D). (c) and (d) are the higher magnification views of boxed areas in A and B. (E) Immunofluorescence image of Col22a1-EGFP forelimb sagittal sections at P0. Blue signals are nuclei stained with DAPI. (e’) and (e’’) are the higher magnification views of boxed areas in E, from MTJ-proximal (e’) and MTJ-distant (e’’) regions, respectively. Dotted lines outline myonuclei, and arrowheads indicate GM130-positive Golgi apparatus. Scale bar: 10 µm (F, G) Rose plots showing the distribution of Golgi apparatus relative to the nucleus for MTJ-proximal (F) and MTJ-distant (G) myonuclei. Bars indicate the percentage of Golgi apparatus in each 10° bin, and dots on the outer ring indicate individual Golgi apparatus.

### Muscle-tendon interface exhibited unexpectedly complex 3D structure with a specialized cell population

Next, we examined MTJ distribution at the tissue level, especially in relation to tendon structure. Tissue clearing with COL1 immunostaining revealed that the COL1+ root-like protrusions with associated MTJs are found at the terminal region of tendon body in SOL and EDL muscles (Fig. 5A-C, Fig. S4A, B). Section immunofluorescence also identified a complex morphology with associated MTJs at the terminus of EDL distal tendon (Fig. S4C). Similarly, COL1+ branch-like protrusions were also identified in the middle part of the tendon body, connecting the tendon body and MTJs embedded within myofibers (Fig. 5D, E). Interestingly, these root-like structures at the terminus or branch-like protrusions were not observed in newborn mice (Fig. S5). These data suggest that the tendon structure and 3D distribution of MTJs changed dynamically during the postnatal period, resulting in a highly interlocked morphology at the muscle-tendon interface.

**Figure 5.**
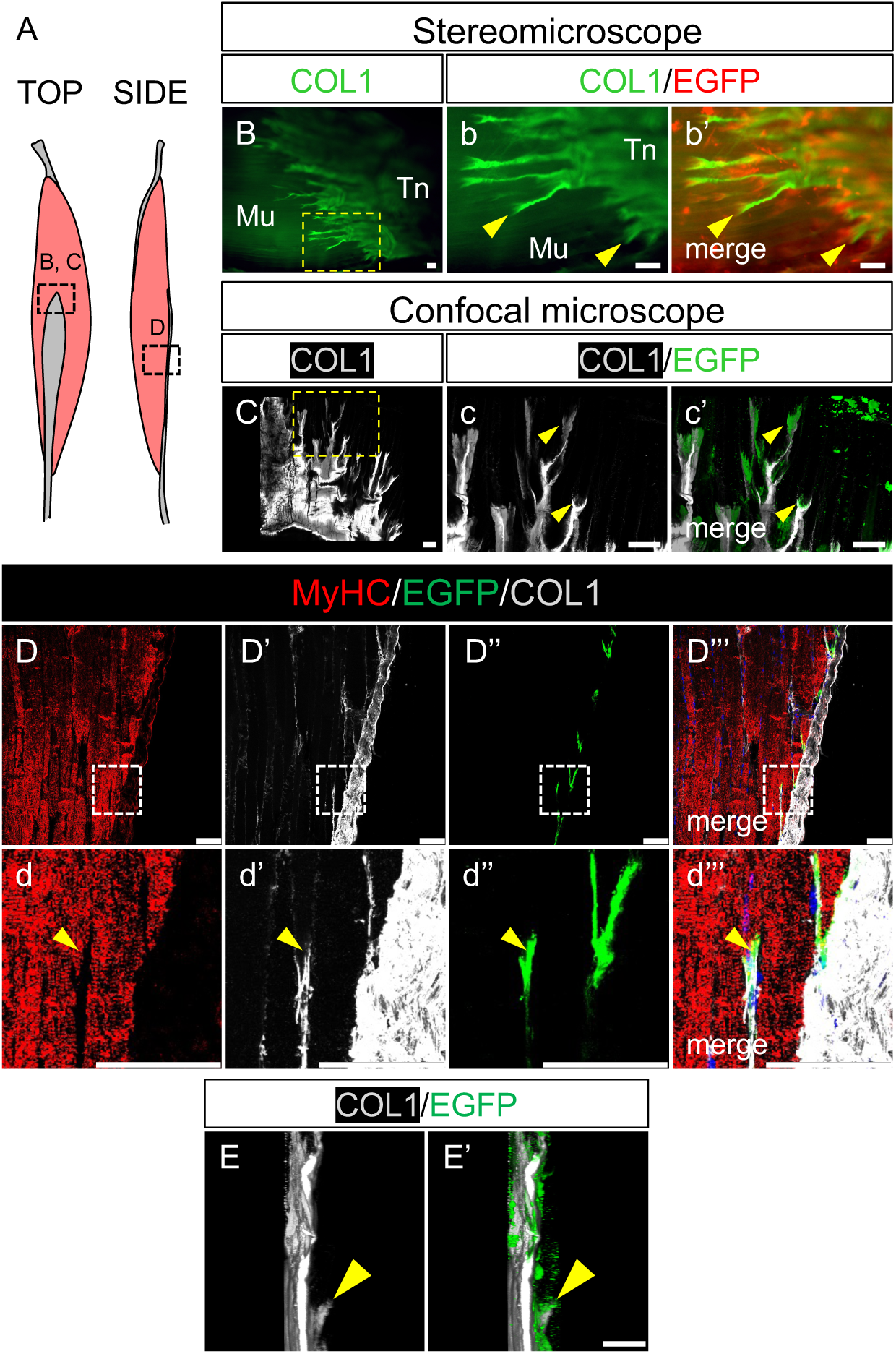
3D structure of muscle-tendon interface. (A) A schematic drawing showing muscle-tendon interface of EDL/SOL muscle and its distal tendon. Top and side views are shown. Boxes indicate the area analyzed in B, C and D, E. (B, C) Immunofluorescence images of tissue-cleared SOL muscle (terminal part) of Col22a1-EGFP knock-in mice. (b) and (c) are higher magnification views of boxed areas in B and C. The yellow arrowheads indicate MTJ distant from the tendon body. Scale bar: 50 μm (D) Immunofluorescence images of EDL muscle (middle part) section of Col22a1-EGFP knock-in mice. (d) is higher magnification views of boxed areas in D. The yellow arrowheads indicate MTJ distant from the tendon body. Blue signals are nuclei stained with DAPI. Scale bar: 50 µm (E) Immunofluorescence images of tissue-cleared EDL muscle (middle part) of Col22a1-EGFP knock-in mice. The yellow arrowheads indicate MTJ distant from the tendon body and COL1 positive protrusion connecting the tendon body. Scale bar: 100 µm

Furthermore, stereomicroscope observation of *ScxCreERT2;Rosa26-tdTomato*;*Col22a1-EGFP* double fluorescent reporter identified that, while the majority of tdTomato+ tenocytes embedded in the tendon body had elongated morphology, a small subpopulation of tenocytes with round morphology was closely associated with MTJs (EDL, Fig. 6A-C; SOL, Fig. S6A-C, magenta and yellow arrowheads). Section analysis confirmed that tdTomato+ *Scx*-lineage cells colocalized with COL22A1-EGFP signal at the tip of myofibers (EDL, Fig. 6D, SOL, Fig. S6D, yellow arrowheads). The morphology of these cells, as well as their nuclei, was roundish (or laterally expanded) and clearly distinct from the elongated tenocytes within the tendon body (Fig. 6D, Fig. S6D, yellow vs magenta arrowhead). Moreover, we found that these cells expressed *Col22a1* mRNA, which is not expressed in normal tenocytes (Fig. 6E, F). In sum, we identified a subpopulation of tenocytes located at the MTJ with specialized morphology and gene expression, potentially contributing to the ECM environment of MTJs as well as branch/root structure formation.

**Figure 6.**
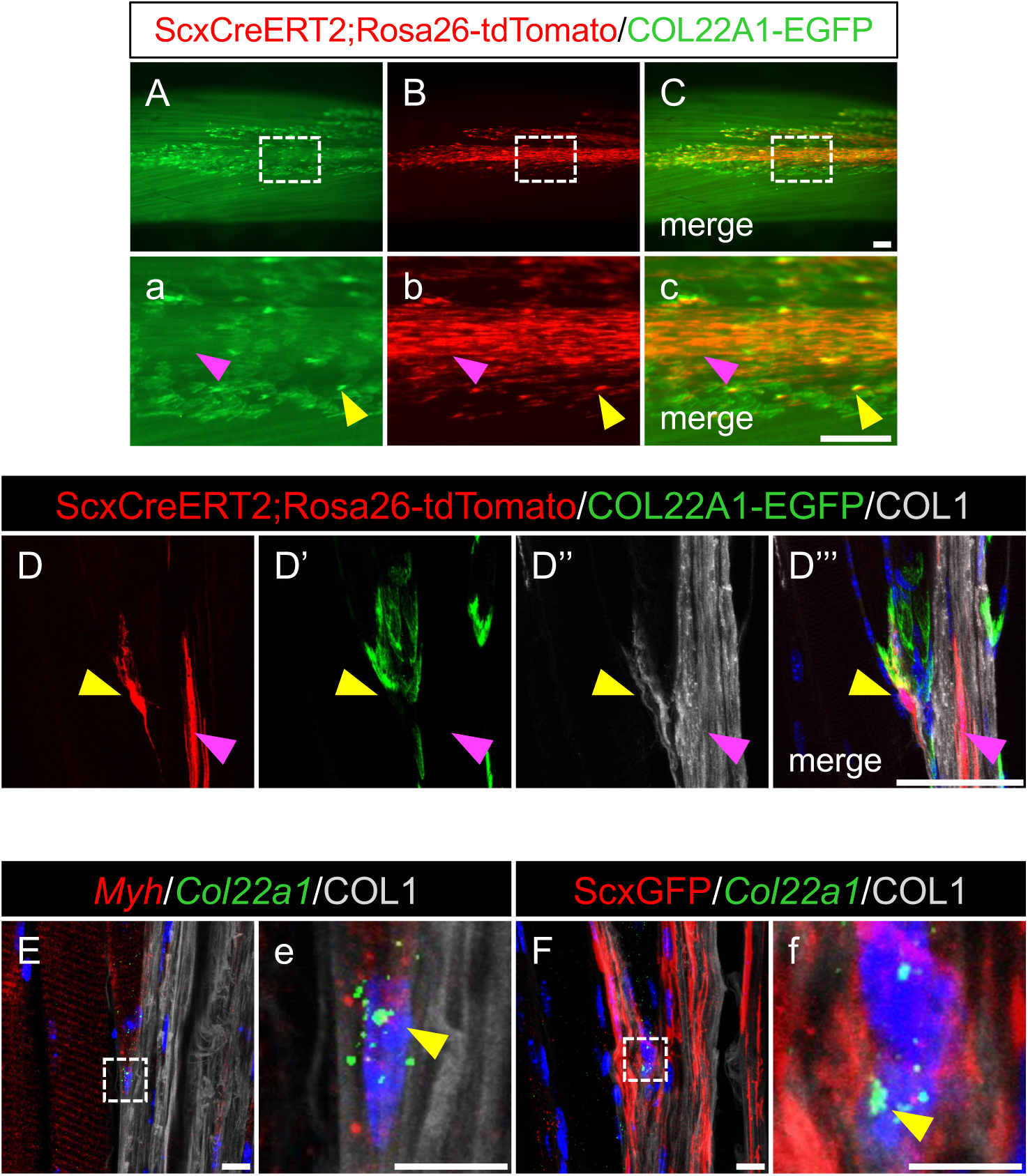
Tenocyte characterization at the muscle-tendon interface. (A-C) Stereomicroscope image of EDL muscle of *ScxCreERT2;Rosa26-tdTomato;Col22a1-EGFP* knock-in mice. (a)-(c) are the higher magnification images of the boxed areas shown in A-C. The magenta and yellow arrowheads indicate tdTomato single positive and tdTomato/EGFP double positive area, respectively. Scale bar: 1 mm. (D) Immunofluorescence images of EDL muscle section of *ScxCreERT2;Rosa26-tdTomato;Col22a1-EGFP* knock-in mice. The magenta and yellow arrowheads indicate tdTomato single-positive and tdTomato/EGFP double positive, and EGFP single positive area, respectively. Blue signals are nuclei stained with DAPI. Scale bar: 20 μm (E) Fluorescent ISH image of EDL muscle of wild-type mice. (e) is the higher magnification view of the boxed area shown in E. The yellow arrowhead indicates *Col22a1* positive nucleus within the COL1 positive area near the tip of *Myh3* positive myofiber. Scale bar: 10 µm (F) Fluorescent ISH and immunofluorescence image of EDL muscle of *ScxGFP* Tg mice. (f) is the higher magnification view of the boxed area shown in F. The yellow arrowhead indicates *Col22a1* positive nucleus within the ScxGFP positive area near the tip of myofiber. Blue signals are nuclei stained with DAPI. Scale bar: 10 µm

### Col22a1-EGFP signal adapts to mechanical loading

Finally, we examined whether the COL22A1-EGFP signal reflects the dynamic adaptation of MTJ for the mechanical loadings. To this end, we applied sciatic denervation as an unloading and tenotomy treatment as an overloading model for hindlimb muscles. As a result, COL22A1-EGFP positive area was significantly increased in the plantaris muscles in tenotomy treated mice, while mean signal intensity remained at a similar level (Fig. 7A-F). In contrast, COL22A1-EGFP signal intensity of EDL muscle in denervated mice was significantly reduced, while signal positive area was unchanged (Fig. 7G-L). Unloading and overloading affected different parameters of the COL22A1-EGFP signal (intensity and area, respectively). This may reflect the complexity of tissue remodeling, in which macroscopic changes in muscle volume and microscopic changes in MTJ structure proceed in parallel. Nevertheless, these results indicate that Col22a1-EGFP reporter signal could be used as a quantitative readout of MTJ adaptation to altered mechanical loading in skeletal muscles.

**Figure 7.**
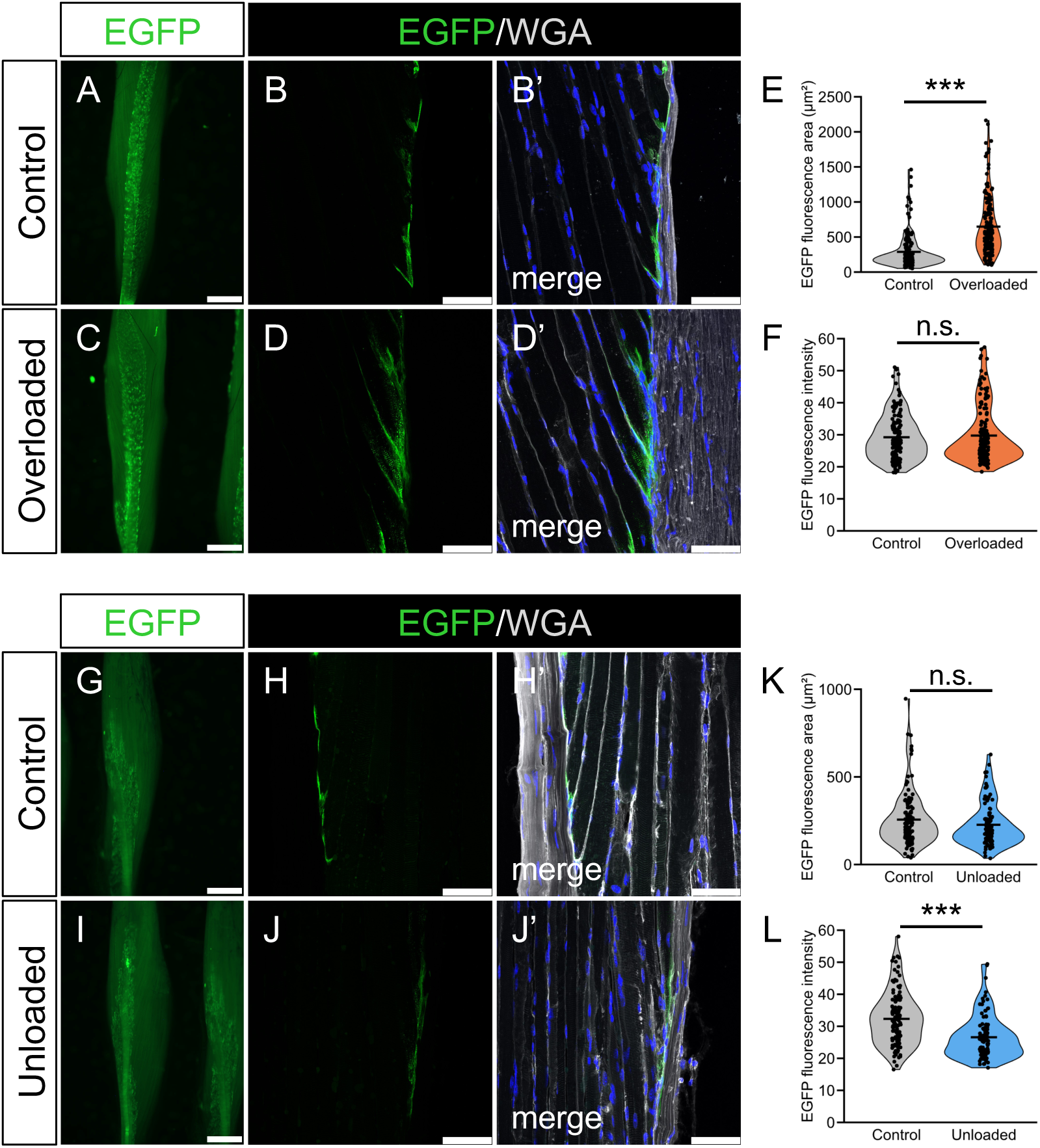
Col22a1-EGFP visualizes MTJ adaptation to altered mechanical loading. (A, C) Fluorescent stereomicroscope image of plantaris muscle of Col22a1-EGFP knock-in mice 14 days after tenotomy operation. “Control” indicates the non-operated side and “Overloaded” indicates the operated side. (B, D) Fluorescence images of plantaris muscle sections shown in A and C. Blue signals are nuclei stained with DAPI. Scale bar: 50 μm. (E, F) The quantification of EGFP signal positive area (E) and mean signal intensity (F). (G, I) Fluorescent stereomicroscope image of EDL muscle of Col22a1-EGFP knock-in mice 14 days after sciatic nerve dissection. “Control” indicates the non-operated side and “Unloaded” indicates the operated side. (H, J) Fluorescence images of EDL muscle section shown in G and I. Blue signals are nuclei stained with DAPI. Scale bar: 50 μm. (K, L) The quantification of EGFP signal positive area (K) and mean signal intensity (L). Statistical analyses were performed using Welch’s t-test, ***p<0.001, n.s., not significant

## Discussion

Historically, MTJs have been studied primarily using fixed tissue sections and immunostaining. Recent studies demonstrated 3D structures of a single MTJ or several MTJs at the tip of isolated myofibers (Hoegsbjerg et al., 2025; Knudsen et al., 2015; Schedel et al., 2026), and our mouse model extends the range and approaches of MTJ analysis, including assessment of tissue-level 3D distribution and dynamic morphological changes. Technically, similar fusion knock-in approaches have been reported for other ECM proteins including COL4A1, COL4A2 and LAMB1 (Jones et al., 2023; Morgner et al., 2023; Wuergezhen et al., 2024). Of note, some fusion knock-ins resulted in loss of function of the targeted protein, and homozygous mice displayed abnormal phenotypes, while others did not. Homozygous Col22a1-EGFP knock-in mice are healthy and fertile and do not show apparent trabecular bone phenotype, indicating that COL22A1-EGFP retains its function, at least in bone. As the loss of Col22 function leads to reduced MTJ function in zebrafish (Charvet et al., 2013), further evaluation is needed for the MTJ of homozygous Col22a1-EGFP knock-in mice.

Our findings revealed the unprecedented variety and complexity of MTJ morphology at both the cellular level (e.g. fiber types) and the tissue level (e.g. 3D distribution). Variation in MTJ morphology among skeletal muscle types could be the result of distinct transcriptional regulations and/or the different mechanical loadings among muscle types. While the MTJ adaptation to the unloading/overloading pointed to the role of mechanical cues (Rocha-Braga et al., 2025), the fiber type dependent variation within a single muscle (soleus) suggests the involvement of intrinsic regulation (Fig. 2). Moreover, the number of *Scx*-lineage cells found at MTJ varied between EDL and SOL muscle, indicating that not only myofiber but also tenocytes could contribute to distinct MTJ morphology. Interestingly, Hoegsbjerg et al reported the fiber type dependent variation of MTJ morphology and associated tendon cells also in human samples (Hoegsbjerg et al., 2025). Additional mechanical models, transcriptional fiber type switching, and further characterization of *Col22a1*+ tenocytes would help elucidate the mechanism underlying MTJ variation (Kurjan et al., 2026; Sadaki et al., 2023; Sadaki et al., 2025). In addition, dynamic postnatal change in 3D distribution of MTJs at the interface of muscle and tendon, including the branch-like protrusions, implies that there could be an adaptation to the allometric growth of tightly connected tissues. MTJ morphology and 3D distribution likely impact the direction and efficiency of force transmission in each muscle type, which will be an interesting topic of future studies.

Embryonic muscle-tendon interaction as well as the MTJ formation process remain largely unclear. COL22A1-EGFP signal was first detected as intracellular puncta within myofibers at E12.5, whereas extracellular signal at the myofiber tip became evident at E14.5. The latter is consistent with previous studies that detected COL22A1 deposition in decellularized forelimbs from E13.5–E14.5 (Leng et al., 2025; Lipp et al., 2023). Because decellularization removes intracellular proteins, our reporter uniquely captures the pre-secretory phase of COL22A1 production, which precedes its extracellular deposition. Detection of earliest COL22A1 expression coincides with the embryonic stage of muscle-tendon connection (Huang et al., 2015; Ono et al., 2023; Sagasaki et al., 2026), indicating that the Col22a1-EGFP could be used as a surrogate marker to detect where and when muscle-tendon connection is formed. As mechanical loading plays an important role in MTJ formation (Lipp et al., 2023; Subramanian et al., 2018), it will be interesting to examine if the Col22a1-EGFP expression or localization changes under conditions where muscle contraction is impaired. Cellular mechanism to transfer and anchor MTJ specific proteins at the myofiber tip is also elusive. Punctate signals of Col22a1-EGFP in embryonic myofiber and biased localization of the perinuclear Golgi apparatus imply that MTJ proteins are transported with polarity toward the myofiber tip. Co-staining of Col22a1-EGFP dots with the markers of transporting vesicles, as well as their live imaging, will provide better understanding of this process. Recently, disorganized localization of COL22A1 protein was reported in *Lama2* mutant mice, indicating that COL22A1 protein could be retained at the MTJ region by forming complexes with LAMA2 and other proteins (Schedel et al., 2026). In this study, we could detect Col22a1-EGFP signals from un-fixed tissue sections, supporting the idea that Col22a1 protein exists as a protein complex rather than in a freely diffusing state in the MTJ region. This suggests that analyzing COL22A1-EGFP interacting proteins could identify local protein networks in an in vivo MTJ environment. MTJ localized protein translation regulation could also be involved (Neves et al., 2026). Further analysis is required to understand the mechanism of MTJ formation and maintenance from a protein localization perspective.

The MTJ is susceptible to injury, especially with high muscle loading of athletes (Edouard et al., 2023; Højfeldt et al., 2025; Jakobsen and Krogsgaard, 2021). A simple method to estimate the function of MTJ could be beneficial for predicting as well as preventing the risk of MTJ injury. Our study demonstrated that COL22A1-EGFP signals reflect the adaptation of MTJ for the change of mechanical loading, implying that Col22a1 expression level could be used as a simple surrogate marker for MTJ status. In line with our findings, *COL22A1* eQTL is related to the athlete injury risk (Miyamoto-Mikami et al., 2020). As such, our reporter mouse model would be a useful tool to evaluate impaired MTJ function upon injury, aging, or disease, as well as its recovery following treatment including mechanical loading.

In summary, Col22a1-EGFP mice revealed unprecedented diversity and dynamic changes of MTJ morphology at the muscle-tendon interface. We believe that the easy visibility, quantitative signal and adoptability of this reporter will open the way to evaluate the MTJ status under various conditions and may help identify the potential interventions to strengthen MTJs and prevent the injury in future studies.

## Acknowledgements

This research is supported by MEXT KAKENHI (22H02636, 24K22257, 26K02842), Terumo foundation, AdAMs for M.I.

## Author Contributions

Y.Y., Y.K., R.S., S.Y., S.J., M.M., K.A., K.O., T.H., H.S., C.S. and S.M. performed the experiments. M.I. planned the study and wrote the manuscript.

## Declaration of Interests

The authors declare no conflicts of interest in association with the present study.

## Notes

### Competing Interest Statement

The authors have declared no competing interest.

### Summary of Updates

Title and abstract is updated to align with the scope of the journal to be submitted (no change in data and results). The device used for micro-CT analysis in Method section has been corrected.

