## Supplementary Files for "Myotendinous junctions form dynamic three-dimensional architectures during development and growth"

Yamasaki, Kawa et al. Figure S1

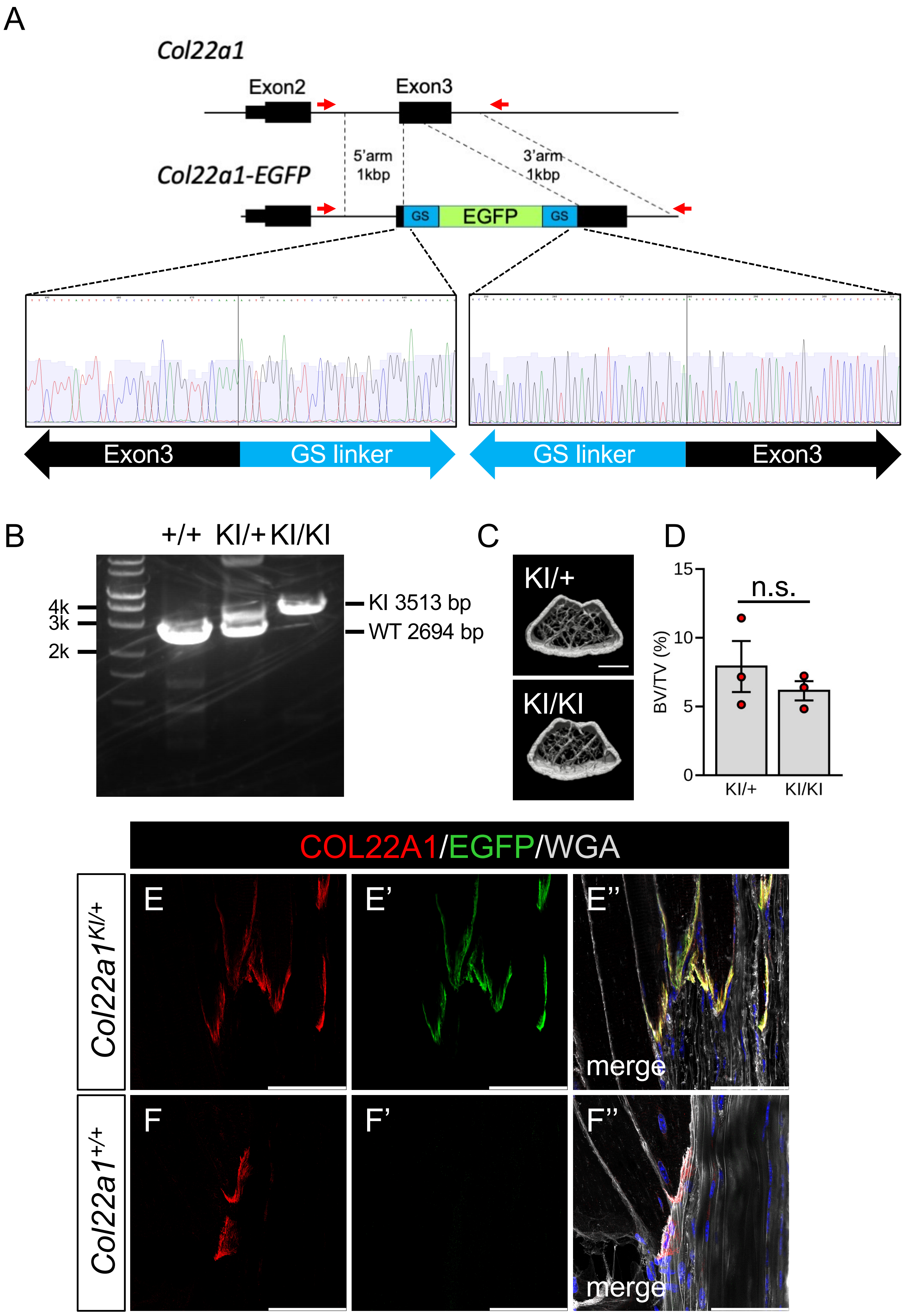

Yamasaki, Kawa et al. Figure S2

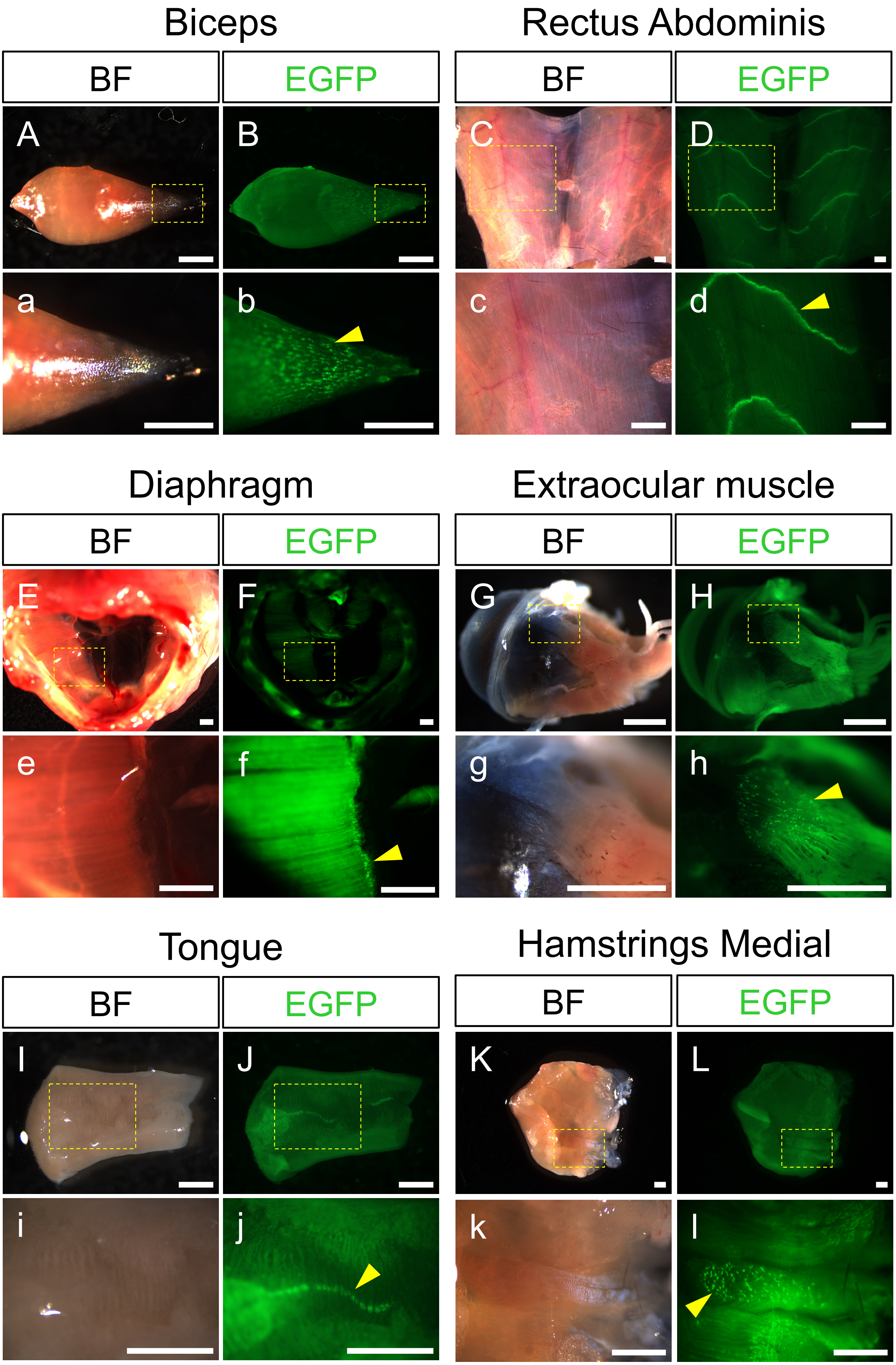

Yamasaki, Kawa et al. Figure S3

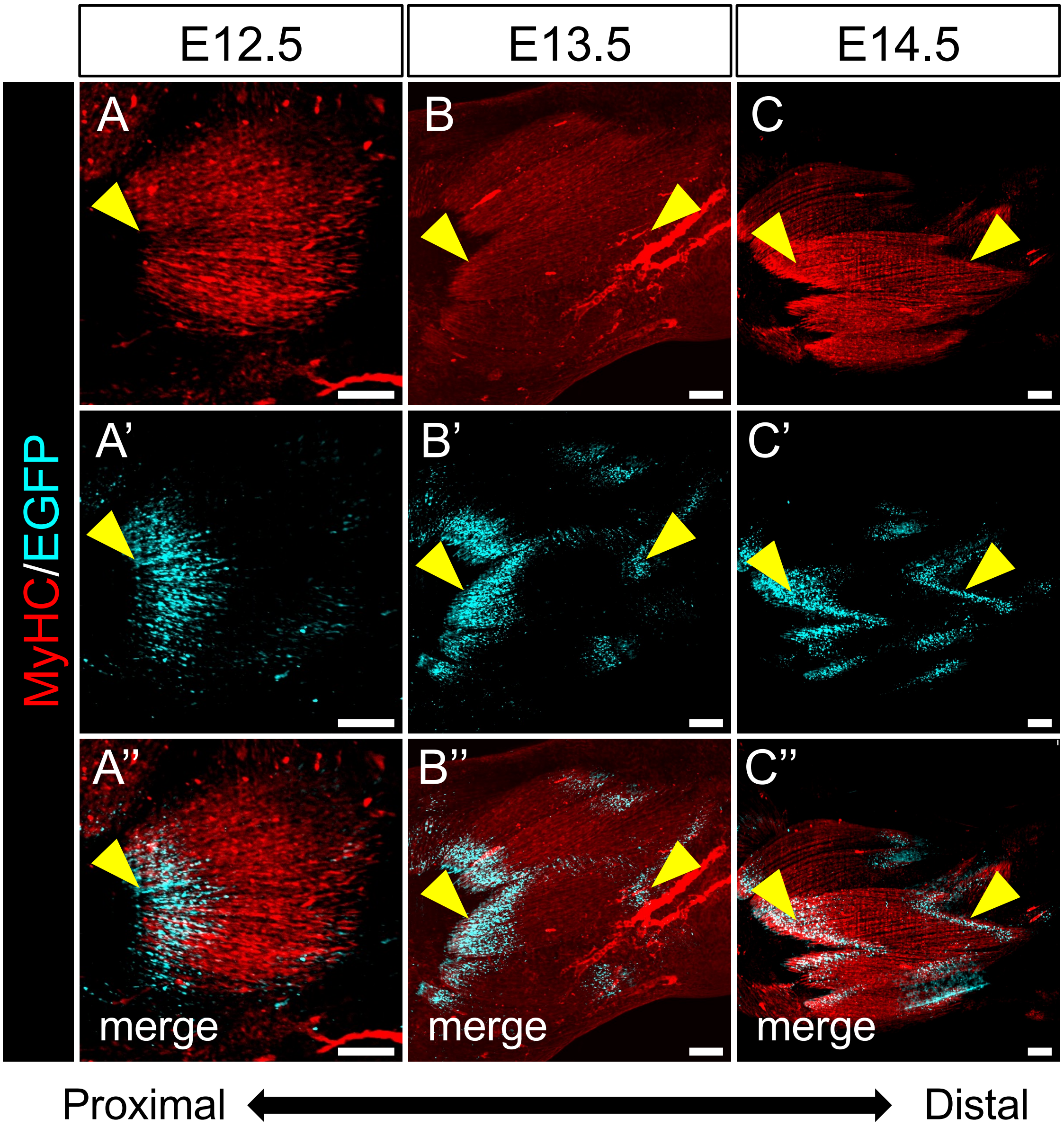

Yamasaki, Kawa et al. Figure S4

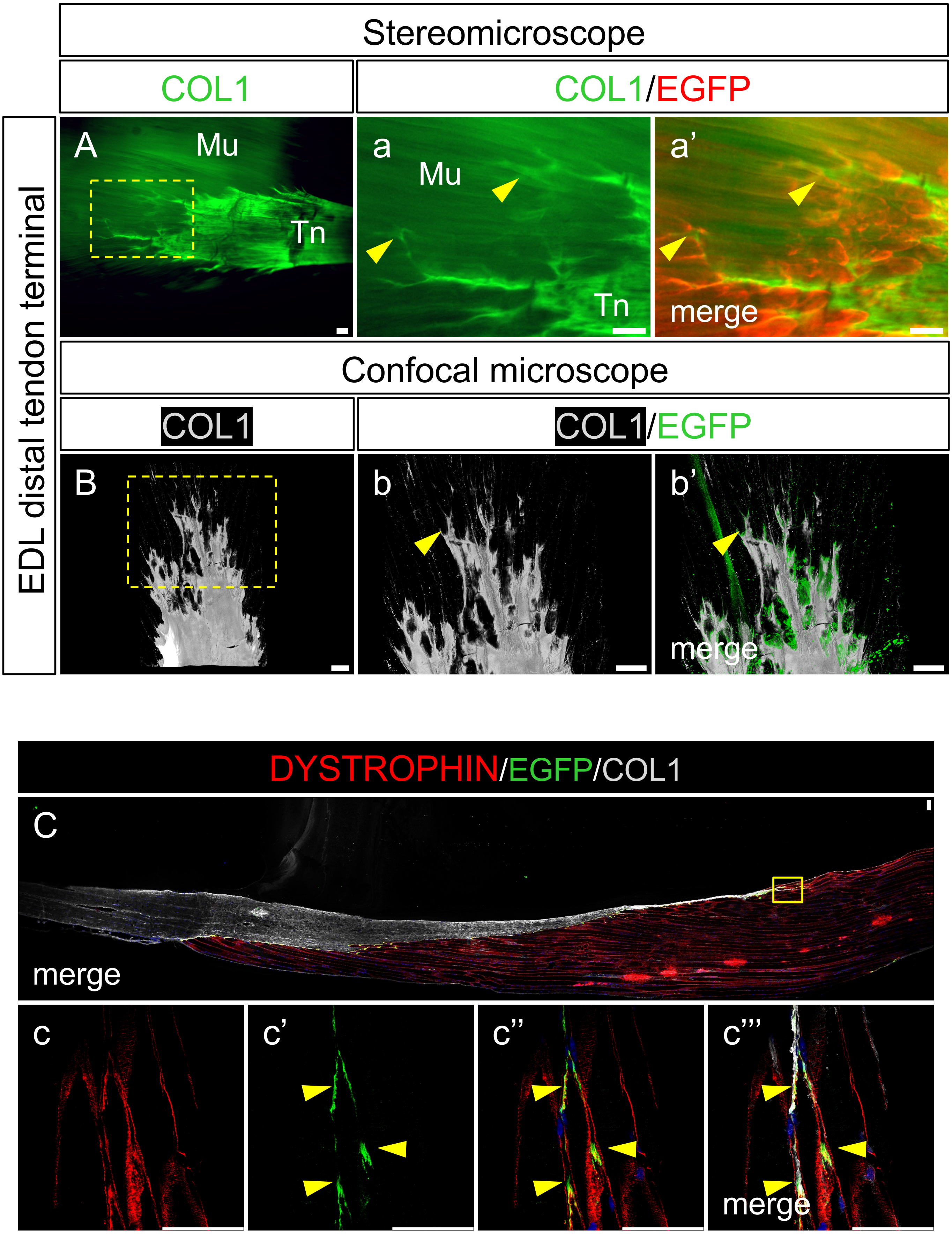

Yamasaki, Kawa et al. Figure S5

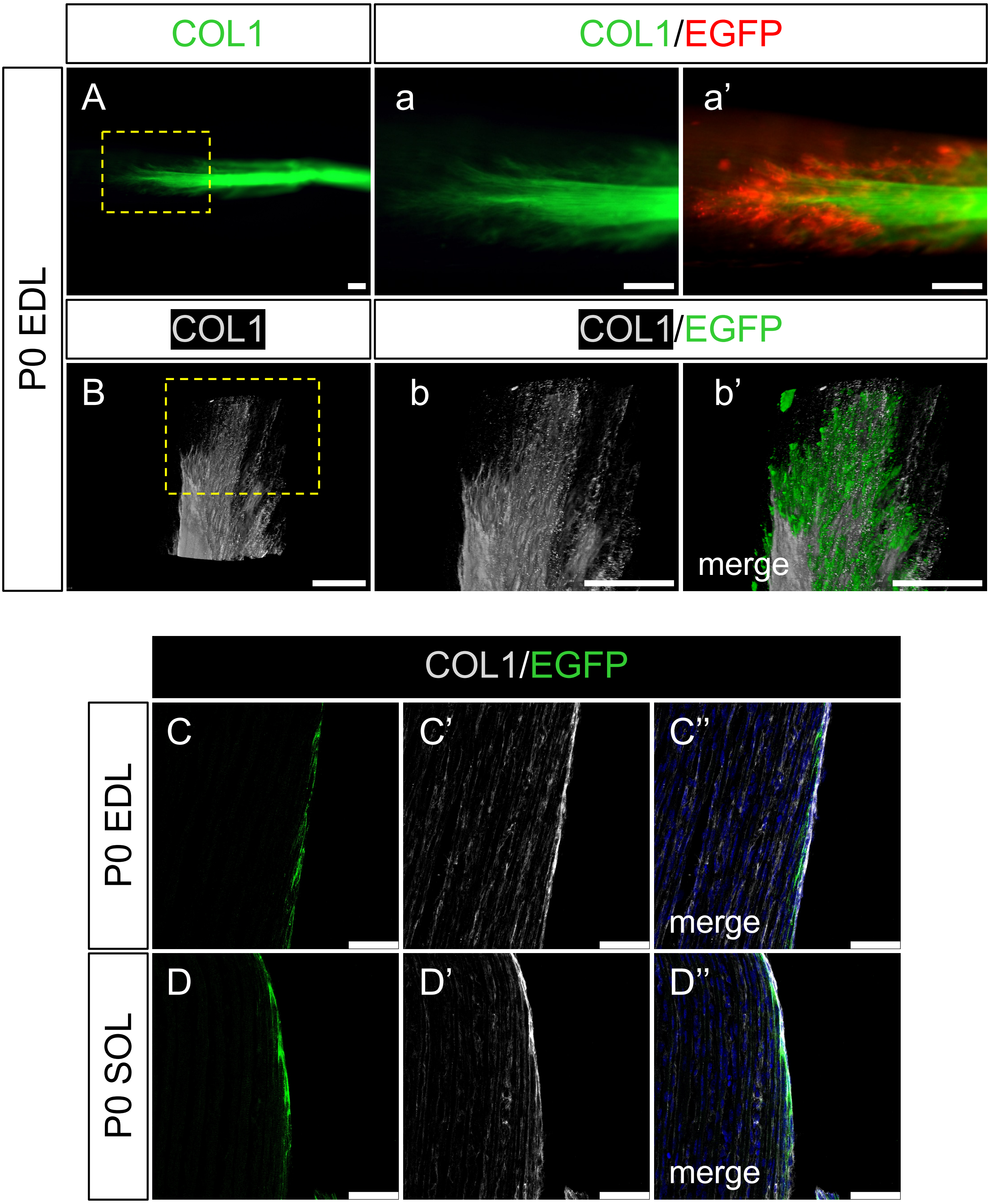

Yamasaki, Kawa et al. Figure S6

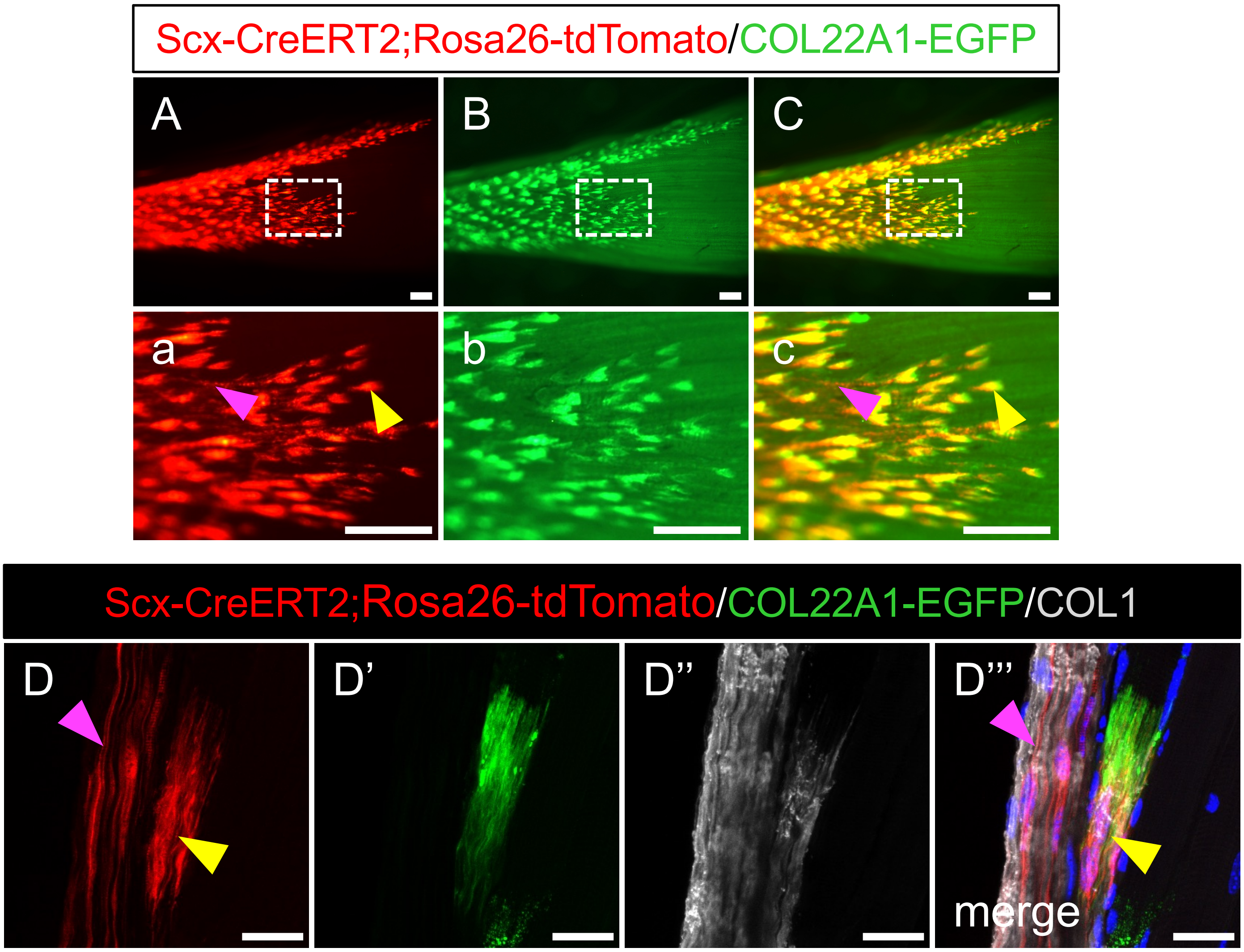

Yamasaki, Kawa et al. Figure S7

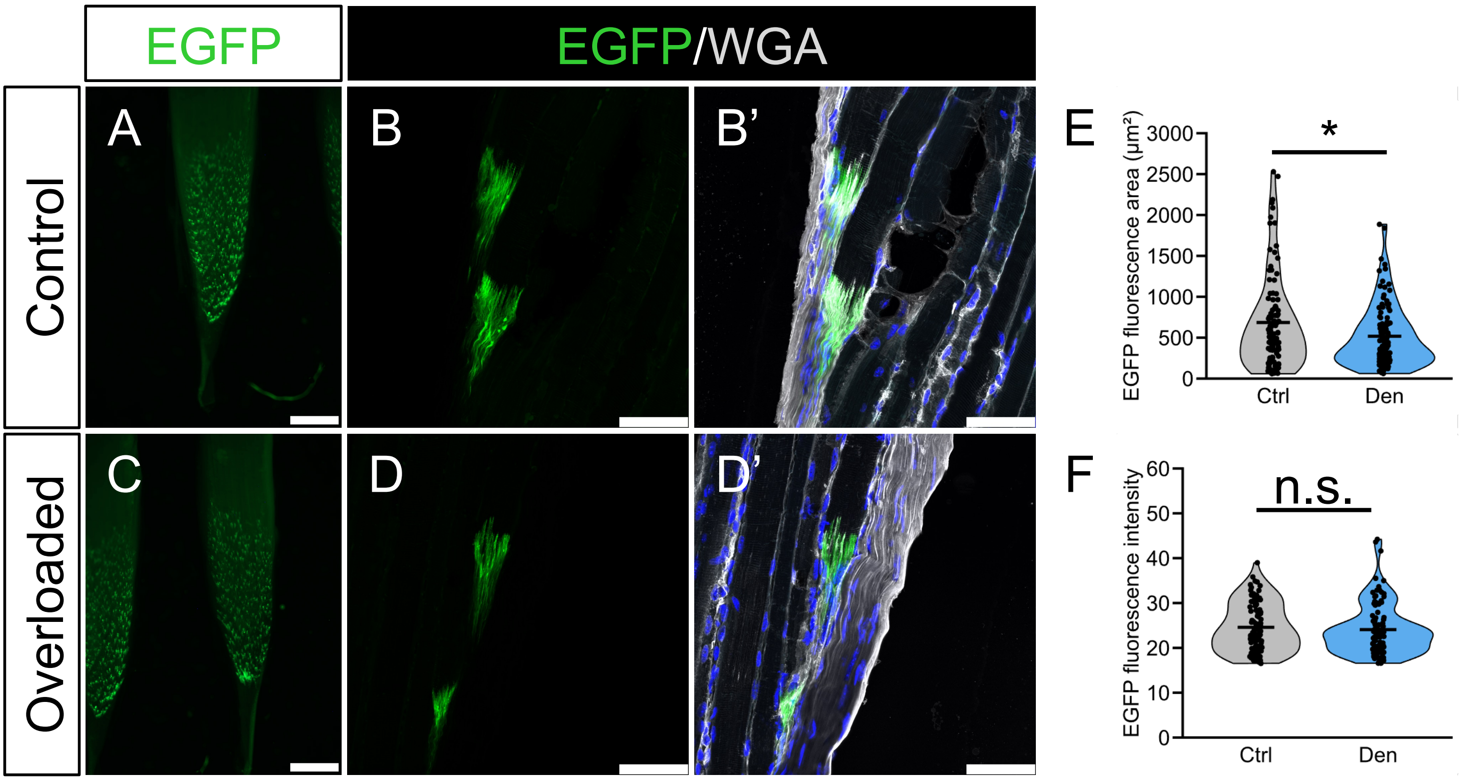

**Supplementary Figure legends**

**Supplementary Figure S1**

(A) A schematic drawing of Col22a1-EGFP knock-in design with the electropherograms showing the boundaries between exon 3 sequence and GS-linker sequence. Red arrows indicate the position of primers used in the genotyping PCR. (B) PCR amplification by the primers indicated in (A) from wildtype (+/+), heterozygous (KI/+), and homozygous knock-in (KI/KI) animals. (C) Representative μCT reconstruction image of the distal femora of 24-weeks-old male Col22a1-EGFP mice. Scale bar: 600 μm (D) μCT-based quantification of the bone volume/tissue volume (BV/TV) of distal metaphyseal trabecular bone shown in (C). Data is shown as mean+/-s.d. and statistical analysis was performed using Welch's t-test, n.s., not significant (n = 3 per group). (E, F) Immunofluorescence images of tibialis anterior muscle section of wild-type ($Col22a1^{+/+}$) and heterozygous Col22a1-EGFP knock-in ($Col22a1^{KI/+}$) mice. WGA, wheat germ agglutinin. Blue signals are nuclei stained with DAPI. Scale bar: 50 μm.

**Supplementary Figure S2**

(A, B) Stereomicroscope image of Biceps muscle of Col22a1-EGFP knock-in mice. (a) and (b) are the higher magnification images of boxes shown in A and B. The yellow arrowhead indicates COL22A1-EGFP signal. (C, D) Stereomicroscope image of Rectus Abdominis muscle of Col22a1-EGFP knock-in mice. (c) and (d) are the higher magnification images of boxes shown in C and D. The yellow arrowhead indicates COL22A1-EGFP signal. (E, F) Stereomicroscope image of diaphragm of Col22a1-EGFP knock-in mice. (e) and (f) are the higher magnification images of boxes shown in E and F. The yellow arrowhead indicates COL22A1-EGFP signal. (G, H) Stereomicroscope image of medial side of Extraocular muscle of Col22a1-EGFP knock-in mice. (g) and (h) are the higher magnification images of boxes shown in G and H. The yellow arrowhead indicates COL22A1-EGFP signal. (I, J) Stereomicroscope image of tongue of heterozygous Col22a1-EGFP knock-in mice. (i) and (j) are the higher magnification images of boxes shown in I and J. The yellow arrowhead indicates COL22A1-EGFP signal. (K, L) Stereomicroscope image of medial side of hamstrings of Col22a1-EGFP knock-in mice. (k) and (l) are the higher magnification images of boxes shown in K and L. The yellow arrowhead indicates COL22A1-EGFP signal. Scale bar: 1 mm.

**Supplementary Figure S3**

(A-C) Maximum intensity projection of light-sheet images of tissue-cleared and immunostained embryonic forelimbs at E12.5 (A), E13.5 (B), and E14.5 (C). The yellow and white arrowheads indicate the proximal and distal end of extensor digitorum communis (EDC) muscles, respectively. Scale bar: 100 µm

**Supplementary Figure S4**

(A, B) Immunofluorescence images of tissue-cleared EDL muscle (terminal part) of Col22a1-EGFP knock-in mice. (a) and (b) are higher magnified views of boxed areas in A and B. The yellow arrowheads indicate MTJs at the tip of root-like structure. Scale bar: 50 µm (C) Immunofluorescence images of EDL muscle section of Col22a1-EGFP knock-in mice. (c) is higher magnified views of boxed areas in C. The yellow arrowheads indicate MTJ at the tip of root-like structure. Blue signals are nuclei stained with DAPI. Scale bar: 50 µm

**Supplementary Figure S5**

(A, B) Immunofluorescence images of tissue-cleared neonatal (P0) EDL muscle (terminal part) of Col22a1-EGFP knock-in mice. (a) and (b) are higher magnified views of boxed areas in A and B. Scale bar: 50 µm (C, D) Immunofluorescence images of neonatal (P0) EDL (C) and SOL (D) muscle section of Col22a1-EGFP knock-in mice. Blue signals are nuclei stained with DAPI. Scale bar: 50 µm

**Supplementary Figure S6**

(A-C) Stereomicroscope image of SOL muscle of ScxCreERT2;Rosa26-tdTomato;Col22a1-EGFP knock-in mice. (a)-(c) are the higher magnification images of the boxed areas shown in A-C. The magenta and yellow arrowheads indicate tdTomato single positive and tdTomato/EGFP double positive area, respectively. Scale bar: 1mm. (D) Immunofluorescence images of SOL muscle section of ScxCreERT2;Rosa26-tdTomato;Col22a1-EGFP knock-in mice. The magenta and yellow arrowheads indicate tdTomato single positive and tdTomato/EGFP double positive area, respectively. Blue signals are nuclei stained with DAPI. Scale bar: 20 µm

**Supplementary Figure S7**

(A, C) Stereomicroscope image of SOL muscle of Col22a1-EGFP knock-in mice 14 days after sciatic nerve dissection. “Control” indicates the non-operated side and “Unloaded” indicates the operated side. (B, D) Immunofluorescence images of SOL muscle section shown in A and C. Blue signals are nuclei stained with DAPI. Scale bar: 50 µm. (E, F) The quantification of EGFP signal positive area (E) and mean signal intensity (F). Statistical analysis were performed using Welch's t-test, * $p<0.05$, n.s., not significant

**Supplemental Table 1**

| Primer name | Primer sequence (5' to 3') |
|---|---|
| Forward primer | CTCTTCTGGCTCCTGTGGTG |
| Reverse primer | CGTTGTCCAATACCCAGGCT |

**Supplemental Table 2**

| Probe name | Probe sequence (5' to 3') |
|---|---|
| Myh-1-P1-S45 | CCTCCACGTaaCTTCCATCTCGGTGTCGCTACTCAT |
| Myh-1-P2-S45 | GGAGGAAAGGTGCAGCTATGCCAAAaaTCCATCTAAGCT |
| Myh-209-P1-S45 | CCTCCACGTaaTACACGTCCTCTGGCTTAACCACCA |
| Myh-209-P2-S45 | TTGTCAAACTTGGGTGGGTTCATGGaaTCCATCTAAGCT |
| Myh-311-P1-S45 | CCTCCACGTaaGACGTGTATCGGTCCTTGAGGTTGT |
| Myh-311-P2-S45 | AGGCCTGAATAGGTGTAGATCATCCaaTCCATCTAAGCT |
| Myh-710-P1-S45 | CCTCCACGTaaAAGCGAGACGAGTTGTCGTTCCTCA |
| Myh-710-P2-S45 | CCAAAATGGATGCGGATGAACTTGCaaTCCATCTAAGCT |
| Myh-887-P1-S45 | CCTCCACGTaaATCAGCAGCAGCTCAATCAGCTCAG |
| Myh-887-P2-S45 | AACGGGTAGTCATAAGGGTTGGTCGaaTCCATCTAAGCT |
| Myh-1145-P1-S45 | CCTCCACGTaaCCCATCAAGTAGGCTGTTTTGTCAG |
| Myh-1145-P2-S45 | GCCTTGAGGAGGTCTGAGGAATTCAaaTCCATCTAAGCT |
| Myh-1287-P1-S45 | CCTCCACGTaaGAACAGCTTCTCGTACACGGACTTG |
| Myh-1287-P2-S45 | CTGGTTAATACGAGTGACCATCCACaaTCCATCTAAGCT |
| Myh-1864-P1-S45 | CCTCCACGTaaCATCTGTTGTAGCGAAGGTGGCATA |
| Myh-1864-P2-S45 | TAGCGACTTTCTTCTTCCCACCGTCaaTCCATCTAAGCT |
| Myh-2380-P1-S45 | CCTCCACGTaaATTCCACACGCATGAGAAACCCTCT |
| Myh-2380-P2-S45 | ACTCCCTCCTCTGCATCATCTTCTGaaTCCATCTAAGCT |
| Myh-2538-P1-S45 | CCTCCACGTaaCTTCATGGTGGCCATCTCCTTCTCA |
| Myh-2538-P2-S45 | CTCATCCTTGGTTTTCTGGAACTCCaaTCCATCTAAGCT |
| Myh-2945-P1-S45 | CCTCCACGTaaATAGTTTCATCCAGGCCTGCCAGTT |
| Myh-2945-P2-S45 | GCCTTCTTCTCCCGTGTTAGCTTTGaaTCCATCTAAGCT |
| Myh-3719-P1-S45 | CCTCCACGTaaTTCTCCAGATTGGCCTTGGATTTGG |
| Myh-3719-P2-S45 | AACTGATCCTCTAGAGTTCGGCAGAaaTCCATCTAAGCT |
| Myh-4339-P1-S45 | CCTCCACGTaaCTAGCACCTTGTCAAAGTTCCTCTG |
| Myh-4339-P2-S45 | TCTCCTCACACTTCGTCTTCCACTCaaTCCATCTAAGCT |
| Myh-5511-P1-S45 | CCTCCACGTaaCTCATACTTCCTCAGGCCCTTCACA |
| Myh-5511-P2-S45 | CTGATATGTGAGCTCCTTAACACGCaaTCCATCTAAGCT |
| Myh-5621-P1-S45 | CCTCCACGTaaCTGCCTCTTGTAGGACTTGACTTTC |
| Myh-5621-P2-S45 | ATTGGCTTGTTCATCAGCCTCCTCAaaTCCATCTAAGCT |
| Myh-5726-P1-S45 | CCTCCACGTaaTGCGATTCTGCGATATCAGCCCGTT |
| Myh-5726-P2-S45 | CGGGTCTTTGCCCGCAGTTTATTGAaaTCCATCTAAGCT |
| Col22a1-430-P1-S41 | GCTCGACGTaaCTCTCCTGCTCGTGAAGACAGGCAT |
| Col22a1-430-P2-S41 | AAGGATTCCCTCTGGGGACAGCCATaaTCCTTTGCAACA |
| Col22a1-482-P1-S41 | GCTCGACGTaaAGTAGAATCCACAGAAGACAGGCCA |
| Col22a1-482-P2-S41 | TGGCAGCCACCATCCCCTCCCCAGAaaTCCTTTGCAACA |
| Col22a1-534-P1-S41 | GCTCGACGTaaACTTTTGCAACCTGCACGTTGGGCC |
| Col22a1-534-P2-S41 | CAGGAGGAAGACCAGATCATACTGCaaTCCTTTGCAACA |
| Col22a1-586-P1-S41 | GCTCGACGTaaCCTTGCCCACACTAGAAGACGTGTC |
| Col22a1-586-P2-S41 | CCCACTGGCGGACTTTCTCAAAGTCaaTCCTTTGCAACA |
| Col22a1-710-P1-S41 | GCTCGACGTaaCCCAGTTCAAAGGCTGTAGTGGGCC |
| Col22a1-710-P2-S41 | TTCACCTCCTCGCGAGAGTTGAAGTaaTCCTTTGCAACA |
| Col22a1-774-P1-S41 | GCTCGACGTaaGTTACCGCCATGGTAGGTGATGCGC |
| Col22a1-774-P2-S41 | ATAACGCAGGGCGTCACCCGTGTTCaaTCCTTTGCAACA |
| Col22a1-857-P1-S41 | GCTCGACGTaaAAGGCACGATTCCCAGGTCTGCCGC |
| Col22a1-857-P2-S41 | TCAGTCAACAGGATGGCCACCTGCTaaTCCTTTGCAACA |
| Col22a1-959-P1-S41 | GCTCGACGTaaCCAACAGCAAAAATGCGGATGCCGG |
| Col22a1-959-P2-S41 | AGTTCCTCCTTCAATGCTGCACCCAaaTCCTTTGCAACA |
| Col22a1-1011-P1-S41 | GCTCGACGTaaCTTGGGTTCTGAGGCAATCTCATCC |
| Col22a1-1011-P2-S41 | GTCGGAAACATGGAAGACATGCGCAaaTCCTTTGCAACA |
| Col22a1-1063-P1-S41 | GCTCGACGTaaCACGGATCTTGTCAATGGCATTGAA |
| Col22a1-1063-P2-S41 | TTTCACAAAGCCGACGCCTCAGCTTaaTCCTTTGCAACA |
| Col22a1-1115-P1-S41 | GCTCGACGTaaACCCGGACACTTGGGCAGAGCACAT |
| Col22a1-1115-P2-S41 | TTGGTATGTTTAAACCGATCTCCTTaaTCCTTTGCAACA |
| Col22a1-1167-P1-S41 | GCTCGACGTaaACCTGTAATTTCCTTGGTTCCTCCA |
| Col22a1-1167-P2-S41 | CACACTGAACAAATCCATCAAGTCAaaTCCTTTGCAACA |
| Col22a1-1219-P1-S41 | GCTCGACGTaaTTTCTCTCTTCCCCAAGATTTCCTT |
| Col22a1-1219-P2-S41 | TCCGTACATAGGAACTCTGGGCTCCaaTCCTTTGCAACA |
| Col22a1-1271-P1-S41 | GCTCGACGTaaCTCTGCACCACAGGGAAGGAACCCA |
| Col22a1-1271-P2-S41 | AGACCTTGGGGAAACACATCTTCAGaaTCCTTTGCAACA |
| Col22a1-1323-P1-S41 | GCTCGACGTaaTGTGACAAAGGCATACTCATCAGGA |
| Col22a1-1323-P2-S41 | CCGTGATGTTTTCCGGAACCGGAAAaaTCCTTTGCAACA |
| Col22a1-1375-P1-S41 | GCTCGACGTaaCTTGCCAGATGTACCAATCTTCTTT |
| Col22a1-1375-P2-S41 | CCTGTGGGATGCCATACTGGTCAATaaTCCTTTGCAACA |
| Col22a1-1427-P1-S41 | GCTCGACGTaaTTCTCACCATCCAGCCGGATAGAGA |
| Col22a1-1427-P2-S41 | CCCACAGCGTTGTACTCAACTGCCTaaTCCTTTGCAACA |
| Col22a1-1516-P1-S41 | GCTCGACGTaaGGTCAAAGAGGTCATCAACTCGGGG |
| Col22a1-1516-P2-S41 | TGCTTAGAGCCATCTTGTGCCAGTCaaTCCTTTGCAACA |
| Col22a1-1568-P1-S41 | GCTCGACGTaaTAGAGGGACACATTCTGGGCCTGGA |
| Col22a1-1568-P2-S41 | AGCGTCTGAACCAGCAAACAGTCAAaaTCCTTTGCAACA |
| Col22a1-1620-P1-S41 | GCTCGACGTaaGATATTCTCCCTCTCCTCAATAGGC |
| Col22a1-1620-P2-S41 | GCCAATCACAGTCTTGCCCTGGATGaaTCCTTTGCAACA |
